# Non-canonical ribosome organization enables translation stalling in neuronal RNA granules

**DOI:** 10.64898/2026.07.18.739316

**Authors:** Jingyu Sun, Lily Drever, Laura Bohorquez, Najia Bouaddouch, Jewel T-Y Li, Thibault Legal, Aldo Hernandez-Corchado, Shuaiqi Guo, Khanh Huy Bui, Hamed Najafabadi, Wayne S. Sossin, Joaquin Ortega

**Affiliations:** Department of Anatomy and Cell Biology, McGill University, Montreal, Quebec H3A 0C7, Canada; Centre de Recherche en Biologie Structurale, McGill University, 3649 Promenade Sir William Osler, Montreal, Quebec H3G 0B1, Canada; Department of Neurology and Neurosurgery, Montreal Neurological Institute, McGill University, Montreal, Quebec, H3A 2B4, Canada; Department of Human Genetics, McGill University, Montreal, Quebec, H3A 0C7, Canada; McGill Genome Centre, Montreal, QC, H3A 0G1, Canada

## Abstract

Neuronal outgrowth to support cognitive functions relies on a timely supply of proteins to distal sites in the neuron. To meet this demand, neurons transport mRNAs and ribosomes in large assemblies called “neuronal RNA granules”. Ribosomes that integrate into these granules initiate protein synthesis in the soma, stall the process, and then become packaged into clusters that are transported to distal sites, where protein synthesis can be reactivated. The exact mechanisms and structure of these ribosome clusters remain unknown. We combined cryo-electron microscopy and cryo-electron tomography to examine these dense ribosome clusters purified from rat brains, which are included in neuronal RNA granules. We found that these clusters contain ribosomes stalled at various stages of elongation and that the mRNA sequences where the ribosomes stall facilitate stalling. The ribosome clusters contain multiple polysomal units, and ribosomes from nearby polysomes interact through rRNA expansion segments, only seen before in disomes of inactive or hibernating ribosomes in neurons. The ribosomes in the clusters form unique three-dimensional arrays that evade detection by the ribosome-associated quality-control and NO-GO decay responses, which normally lead to recycling of 40S and 60S subunits, and degradation of the mRNA and arrested nascent proteins. Overall, this study helps uncover specialized mechanisms in neurons that enable local protein translation at distal sites, supporting proper protein homeostasis, both crucial for neurodevelopment and optimal neuronal function.

**KEY POINTS:**

1. Ribosomes in neurons organize into unique clusters that are incorporated into “neuronal RNA granules”.
2. Approximately half of the ribosomes in these clusters are stalled in elongation.
3. The mRNA sequences read by these ribosomes facilitate stalling.
4. Ribosomes within the clusters interact through rRNA expansion segments.
5. Ribosomes in the clusters form unique 3D structures that are not detected by the ribosome-associated quality-control response.

## INTRODUCTION

In most cell types, the cytoplasm surrounds the nucleus. Instead, neurons have a cellular soma containing the nucleus and only a small fraction of the cytoplasm. Approximately 95% of the neuronal cytoplasm extends from the body, forming the axon and thousands of extensions called dendrites. This unique morphology allows neurons to form networks capable of receiving, processing, and transmitting signals. Neuronal extensions contain more than 2,500 different proteins, and their abundance changes with neuronal outgrowth and synaptic plasticity (1,2).

Because the ribosomes and mRNAs needed for protein synthesis are generated in the soma, maintaining and remodeling the proteome across the neuron’s extensive network of processes— often far from the cell body—poses unique challenges. Even using the fastest protein transport mechanism in neurons, it can take hours—and in axons that extend several meters, even days—for a protein to reach the distal end of these projections. However, neurons require responses on timescales of seconds to minutes (3). Consequently, localization of mRNA and ribosomes, and on-site protein synthesis, have evolved in neurons to timely supply new proteins to distal regions (4). This mechanism allows neurons to act as decentralized systems, in which some functions do not entirely depend on the nucleus and soma (5). By synthesizing proteins on-site, neurons can respond dynamically to local signals and support critical processes ranging from synaptic maturation to structural guidance (6). Local translation is particularly high during development, as axons and dendrites actively navigate and form new connections (7). Not surprisingly, disruptions in the localization or translation of these mRNAs are increasingly linked to neurological disorders, highlighting the importance of this process for long-term brain health (8).

On-site protein synthesis in neuronal extensions requires a transport system that delivers mRNAs to the correct location. This system is often mediated by RNA-binding proteins (RBPs) that bind specific mRNAs and regulate their transport and translational status; these RBPs–mRNA complexes are commonly referred to as messenger ribonucleoproteins (mRNPs). In neurons, mRNPs are transported to distal sites either one at a time (9,10) or in granules containing multiple mRNPs that are stalled at translation initiation (11). There are many other types of RNA granules found in most cell types, including processing bodies (P-bodies) (12), stress granules (13), and miRNPs (11). In addition to these structures, neurons also contain specialized transport granules referred to as “neuronal RNA granules”, which are distinguished from other RNA granules by their inclusion of 80S ribosomes (14–19) stalled at the elongation step (20).

The elongation phase of protein synthesis comprises multiple steps, and as the ribosome performs them, its structure undergoes large-scale intra– and inter-subunit rearrangements (21,22). The tRNA molecules also walk the three tRNA binding sites in the intersubunit space during elongation: the aminoacyl (A)-site responsible for binding and decoding incoming aminoacylated tRNAs, the peptidyl (P)-site responsible for orienting the polypeptide-bearing P-site tRNA for efficient transamidation, and the exit (E)-site responsible for subsequent release of deacylated tRNA. An elongation cycle begins with the delivery of the aminoacyl-tRNA (aa-tRNA) to the A-site, complexed with eEF1A-GTP, and involves decoding, GTP hydrolysis, release of eEF1A-GDP, and finally tRNA accommodation. The accommodation step is simultaneous to the 40S subunit rolling motion, which is a ∼ 6° rotation of the 40S subunit toward the uL1 stalk around the long axis of the small subunit. The three tRNA molecules present in the ribosome at this point exist in the classical PRE with A/A-, P/P– and E/E-tRNAs configurations. Subsequently, the peptide bond forms, resulting in a peptidyl– tRNA in the A-site and a deacylated tRNA in the P-site.

To reset the ribosome for the next round of elongation, the ribosome must translocate the tRNA and mRNA molecules by one codon. To this end, the entire 40S subunit rotates by 9° in an anticlockwise direction with respect to the 60S subunit (ratcheting motion), at the same time that the head in the 40S subunit swivels by 13.5° also in an anticlockwise direction. This causes the tRNA in the E/E position to be released and the other two tRNAs to adopt the hybrid A/P and P/E positions (Rotated 1 & 2 states). The rotated translation intermediate is bound by eEF2-GTP, allowing the 40S subunit to back-rotate clockwise, reverting the ratcheting and swivelling motions and allowing the tRNAs and mRNA to move forward. The 40S subunit rolling motion is also reverted at this stage. These motions are also accompanied by the hydrolysis of the GTP molecule and the release of eEF2-GDP. This unrotated translation 80S intermediate is called classical POST state, contains a P/P and an E/E tRNA molecule, and is ready to load a new aa-tRNA-eEF1A-GTP in the A-site to initiate a new elongation cycle.

It remains largely unknown how ribosomes within neuronal RNA granules stall during elongation and how they assemble into ribosome clusters. In this study, we combined cryo-electron microscopy (cryo-EM) and cryo-electron tomography (cryo-ET) to image ribosomes purified from rat brains using sucrose gradient ultracentrifugation to separate these distinct ribosomal clusters. We used image classification approaches to characterize this heterogeneous mixture of ribosomes and found that the ribosomes in the dense clusters from the pellet fraction were largely stalled in specific elongation states. Surprisingly, ribosomes migrating in a sucrose gradient in fractions equivalent in size to the polysome and monosome fractions were also primarily composed of stalled ribosomes, suggesting they are likely an earlier precursor of the dense ribosomal clusters in the pellet. Sequence analysis of the mRNA ribosome-protected fragments (RPFs) indicated that the specific nucleotides at the mRNA exit site and the untranslated nucleotide sequences downstream of the A site, possibly through a secondary structure, such as an RNA G-quadruplex, that impedes ribosome translocation, may play an important role in promoting stalling. Characterization of the higher-order organization of dense ribosome clusters in the pellet fraction revealed that individual clusters typically contain multiple polysomes, disomes, and monosomes. Ribosomes from different polysomes within the same cluster interact via rRNA expansion segments. Neighboring ribosomes on the same mRNA were sometimes spaced far enough apart to avoid direct contact. However, in some cases, they were collided into disomes or longer arrays of ribosomes with 10 or more particles. Notably, these ribosomes interacted through interfaces that differ from those observed in canonical ribosome collisions, which typically trigger the ribosome quality control (RQC) and NO-GO decay responses and lead to recycling of 40S and 60S subunits, degradation of the mRNA and of the arrested nascent proteins (23). These findings provide insights that explain how ribosomes in neuronal RNA granules can evade these stress responses and remain stably stalled.

## MATERIAL AND METHODS

### Purification of ribosome clusters

Neuronal ribosomes were purified from whole-brain homogenate harvested from postnatal day (P5) Sprague Dawley rats of both sexes, using a protocol adapted from a previous study (14). Five P5 rat brains were homogenized in RNA Granule Buffer (20 mM HEPES, pH 7.4; catalog #BP310-500, Thermo Fisher Scientific), 10 mM MgCl_2_ (catalog #M33-500, Thermo Fisher Scientific), 100 mM KCl (catalog #P330-500, Thermo Fisher Scientific) supplemented with 20 mM DTT (catalog #BP172-25, Thermo Fisher Scientific), 0.04 mM spermine (catalog #S2876, Sigma Aldrich), 0.5 mM spermidine (catalog #S2501, Sigma Aldrich) and EDTA-free protease inhibitor (catalog #04693132001, Roche). Homogenate was centrifuged for 15 min in a Thermo Fisher Scientific T865 fixed-angle rotor at 6,117 × g at 4°C to spin down cellular debris. The supernatant was treated with 1% IGEPAL CA-630 (catalog #I8896, Sigma-Aldrich) for 5 min at 4°C on a rocker. The sample was then loaded onto a 2 ml 60% sucrose (catalog #8550, Calbiochem) cushion (dissolved in RNA Granule Buffer) in a Sorvall 36 ml tube (Kendro, catalog #3141, Thermo Fisher Scientific), filled to the top with additional RNA Granule Buffer and centrifuged for 2h in a Thermo Fisher Scientific AH-629 swing-bucket rotor at 56,660 × g at 4°C to achieve the ribosome pellet. The pellet was resuspended in RNA Granule Buffer, gently dounced, and loaded over a 15–60% linear sucrose gradient (gradient was made with RNA Granule Buffer) that was prepared in advance using a gradient maker (Biocomp Gradient Master) and centrifuged for 45 min at 56,660 × g at 4°C in a Thermo Fisher Scientific AH-629 swing bucket rotor. Fractions were then collected from the top, and the pellet fraction was collected by resuspending the pellet at the bottom of the tube with RNA Granule Buffer. In addition, we collected fractions 2/3 and 5/6 for analysis, since they correspond to the fractions in which monosomes and polysomes in other cells migrate. Fractions 2,3,5 and 6 were collected as 3.5 mL fractions, and the others were collected as 3 mL fractions. For the samples purified with cycloheximide treatment, 0.1 mg/mL cycloheximide was added to all buffers during homogenization and purification. In addition, liver polysomes were prepared using 20 livers from the same rats, following the same protocol used for neuronal polysomes without cycloheximide. The only difference was that the supernatant from the first centrifugation run was treated with 1% sodium deoxycholate (DOC) (catalog #D6750, Sigma-Aldrich) instead of 1% IGEPAL CA-630, as this greatly increased the yield of ribosomes from liver.

For the samples used for cryo-EM single-particle analysis, the samples obtained from the last step of purification were treated with 100 U of nuclease (Ambion RNase I, log #AM2294) for 30 min at 4°C. The nuclease was quenched with 100 U of Invitrogen SuperaseIN (log# AM2696, Thermo Fisher Scientific), and the samples were rerun on a fresh 20% sucrose cushion using an AH-629 swing-bucket rotor at 56,660 × g at 4°C for 2hr. A volume of 200 µl of RNA granule buffer was used to resuspend the ribosomal pellets.

### Immunoblotting and quantification of enrichment

For immunoblotting, fractions 2/3 and fractions 5/6 were ethanol precipitated and the pellet was resuspended with RNA granule buffer. We used 1:5 and 1:10 dilutions of the samples for the immunoblots. SDS sample buffer was added to each sample before loading onto a 12% acrylamide gel. The resolved proteins were transferred onto a 0.45mm nitrocellulose membrane (catalog #1620115, Bio-Rad Laboratories) for immunoblotting. Then, the membranes were blocked with 5% bovine serum albumin (BSA) (catalog # A9647, Sigma Aldrich) in Tris-buffered saline with Tween (TBS-T) before incubation with primary antibodies rabbit anti-S6 (1:1000) (catalog #2217, Cell Signaling Technology), rabbit anti-FMRP (1:500)(catalog #4317, Cell Signaling Technology), rabbit anti-UPF1 (1:1000)(catalog #ab133564, Abcam), mouse anti-Stau2 (1:500)(catalog #MM0037-P, MediMabs), rabbit anti-eEF2 (1:1000)(catalog #2332, Cell Signaling Technology), rabbit anti-Pur-alpha (1:1000)(catalog #ab79936, Abcam), rabbit anti-eEF1A (1:1000)(catalog #11402-1-AP, Proteintech), rabbit anti-EBP1 c-terminus (1:1000)(catalog #A02791-2, Boster Bio), mouse anti-hnRNPA2B1 (catalog #NB120-6102, Novus Bio), rabbit anti-TIA1 (catalog #12133-2-AP, Proteintech), mouse anti-G3BP1 (catalog #H00010146-M01, Abnova). Membranes were washed with TBS-T after incubation. Detection was done using HRP-conjugated secondary antibodies anti-rabbit HRP (1:10,000) (catalog #31460, Thermo Fisher Scientific) and anti-mouse HRP (1:10,000)(catalog #31430, Thermo Fisher Scientific). ECL reaction (catalog #NEL105001EA, PerkinElmer) was performed for film imaging, the images were scanned, and signal intensity was quantified by ImageJ software. We selected full-lane ROIs and quantified single bands corresponding to the observed kilodalton size of each protein. For each protein, we calculated enrichment in fractions 2/3 and 5/6 relative to the pellet by dividing fractions 2/3 by the pellet or fractions 5/6 by the pellet. We then used the corresponding anti-S6 signal intensity normalized to the pellet to quantify the amount of protein per S6 ribosomal protein for each fraction. Following this normalization, biological replicates were averaged. To determine the distribution of each RBP across fractions, the percentage of protein in each fraction was calculated using the normalized values. This calculation was performed separately for each protein and each biological replicate. For statistical analysis, a one-way ANOVA was performed for each fraction (Frac 2/3, Frac 5/6, and Pellet) to compare the percentage distribution between proteins. Tukey’s post hoc multiple comparisons tests were then used to identify significant pairwise differences between proteins within each fraction. Graphs were made using R studio.

### Negative-staining electron microscopy

Samples imaged under negative-staining electron microscopy included the pellet fraction, fractions 5/6 and 2/3, all purified from P5 rat brains, and polysomes purified from rat livers. To image these samples, we used copper 400-mesh EM grids coated with a continuous carbon film. For all samples, ribosome concentration was adjusted to ∼32 ng/µl (∼10 nM) in RNA Granule Buffer before applying them to the grids. Before applying the samples, grids were glow-discharged at 15 mA for 15 s. Grids were floated for 2 minutes in a 5 µl drop of sample deposited in a piece of parafilm. Upon incubation, the drop of sample was removed by blotting the grid with filter paper (Whatman #1), and grids were floated in a 5 µl drop of 1% uranyl acetate for 1 minute. The stain was then removed by blotting with filter paper and subsequently allowed to dry in the air. EM Images were acquired on a Tecnai G2 Spirit Twin microscope operating at 120 kV, using a NanoSprint15 MK2 CMOS camera at a nominal magnification of 60,000x. The resulting images were recorded at a defocus of approximately −2.7 μm and had a calibrated pixel size of 1.8 Å/pixel.

### Cryo-EM sample preparation and data collection

The pellet fraction, fractions 5/6 and 2/3, all purified from P5 rat brains were imaged by cryo-EM using the following method. All samples were diluted to 180 nM in RNA Granule Buffer before vitrification using a Vitrobot Mark IV (Thermo Fisher Scientific Inc.). Cryo-EM grids (c-flat CF-2/1-3Cu-T) were washed in chloroform for two hours and glow-discharged in air at 10 mA for 13 seconds. Approximately 3.6 μl of the sample was applied to the grid before plunging into liquid ethane. The Vitrobot parameters for vitrification were set to a blotting time of 3 seconds and a blot force of +1. The Vitrobot chamber was maintained at 25 °C with 100% relative humidity during the process.

All cryo-EM datasets for single-particle analysis were collected using SerialEM software (24,25) in the FEMR-McGill Titan Krios electron microscope operated at 300 kV. Movies were collected using the Gatan K3 direct electron detector mounted on the microscope and equipped with a Quantum LS imaging filter. Data collection parameters for each dataset are described in Supplementary Tables S1 to S6.

### Image processing for single-particle analysis

A similar image processing workflow for single-particle analysis was applied to all datasets using cryoSPARC software versions v4.5 and v4.7 (26). Cryo-EM movies were collected and imported into cryoSPARC. Beam-induced motion was corrected for all movies using Patch Motion Correction with default settings. All frames from each movie were used, with a B-factor of 500 applied, and only information up to 5 Å was considered during alignment. A calibrated smoothing constant of 0.5 was applied to trajectories. CTF parameters were estimated using Patch CTF estimation with default settings. During estimation, the resolution range was set between 4 to 25 Å, and the defocus range between 1,000 to 40,000 Å. The averaged micrographs were subjected to Manually Curate Exposures to exclude micrographs of poor quality.

For particle picking, all micrographs were denoised by the Micrograph Denoiser program using default settings. Then, Blob Picker was first applied to 2,000 micrographs using a circular blob with minimum and maximum particle diameters of 200 Å and 558 Å, respectively. The picked particles were extracted with a box size of 448 pixels, downsampled to 112 pixels and subjected to 2D classification to generate 2D templates for the subsequent Template Picker on the same 2,000 micrographs. In the 2D Classification job, 50 classes were requested, and only representative views of 80S ribosomes were selected as 2D templates for the next step. The Template Picker was run with default settings, and the picked particles were curated again using 2D classification to remove junk particles. The curated particles were used to train a model in the Topaz program (27) on the same 2,000 micrographs, with default settings, including a minibatch size of 128 and an estimated particle diameter of 488 Å. The trained model was then used to pick particles from all micrographs. Particle curation was conducted in two steps. First, two cycles of 2D Classification were performed on all particles obtained from the previous step. Junk particles were discarded, and the good particles were selected and subjected to the next curation phase, which combines Ab-Initio Reconstruction and Heterogeneous Refinement. For the Ab-initio reconstruction job, we requested five classes with a class similarity of 0.1, and the resolution range considered was between 35 and 12 Å. The initial models generated from the Ab-Initio job were subsequently used in the Heterogeneous Refinement using default parameters to separate the particles into multiple 3D classes. The particles assigned to the class with unidentifiable features were discarded, whereas the particles assigned to classes with identifiable features were merged for subsequent processing.

The 3D structural heterogeneity of the particles was explored and separated through three major steps. First, the curated particles were used to generate a consensus map using Non-Uniform Refinement with default settings and C1 symmetry. The aligned particles were then subjected to 3D Variability Analysis, with 3 orthogonal principal modes requested to analyze the heterogeneity, and the resolution filtered at 9 Å before analysis. The cluster mode in the 3D Variability Analysis Display was then used to separate the structural heterogeneity of all particles. The number of requested clusters varied between experiments, ranging from 3 to 5. Two rounds of the 3D Variability Analysis combined with the 3D Variability Analysis Display were performed in this step. The resulting maps from the 3D Variability Analysis Display were visually inspected via UCSF Chimera and Chimera X (28,29), and groups of particles representing similar structural features were merged into classes. In the second step, the 3D Variability Analysis was performed for each class with a focused mask applied to the interface between the 60S and 40S ribosomal subunits. The cluster mode in the 3D Variability Analysis Display was then used to resolve heterogeneity in tRNA conformations. Two rounds of the focused 3D Variability Analysis and 3D Variability Analysis Display were performed, and the groups of particles with identical tRNA states were merged.

To obtain a high-resolution structure for each class, groups of particles representing similar structural features were merged and extracted with a full box size of 448 pixels. The Non-uniform Refinement was performed for each class with C1 symmetry, with options ‘optimize per-particle defocus’, and ‘optimize per-exposure group CTF parameters’ turned on. The ‘Fit Spherical Aberration’, ‘Fit tetrafoil’ and ‘Fit anisotropic Magnification’ were also activated for Global CTF Refinement. The high-resolution refinement maps and particles were subjected to the Reference-Based Motion Correction program to estimate the precise trajectories of each particle. The corrected particles were subsequently used as inputs to rerun the Non-uniform Refinement for each class using the same parameters to obtain the final refinement maps. Average and local resolution estimates were obtained using the gold-standard approach (30). Cryo-EM map visualization was performed in UCSF Chimera and Chimera X (28,29).

### RNA sequencing data analysis

Ribosome profiling libraries were sequenced as three multiplexed samples per lane, distinguished by 4–5 nucleotide inline barcodes (NI810: ATCG, NI811: AGCTA, NI812: CGTAA) ligated upstream of the 3’ adapter. Demultiplexing and adapter trimming were performed simultaneously using Cutadapt v2.10 in single-end mode, no indels permitted, a minimum overlap of 5 nt, and untrimmed reads discarded (31). Reads shorter than 15 nucleotides post-trimming were removed. Five-nucleotide unique molecular identifiers (UMIs) were then extracted from the 5’ end of each demultiplexed read using UMI-tools v1.1.4 with the pattern NNNNN and appended to read headers for downstream duplicate discrimination (32). Ribosomal RNA reads were depleted using BBDuk from the BBMap suite v39.01 via k-mer-based filtering (k = 31) against the ribokmers.fa.gz reference database. Filtered reads were aligned to the Mus musculus GRCm38 transcriptome (Ensembl release 102, including coding and non-coding sequences) using STAR v2.7.10, permitting a maximum of one mismatch and retaining uniquely mapped reads only, with output as coordinate-sorted BAM files (33). PCR duplicates were removed with UMI-tools dedup using the “unique” method, which groups reads by genomic position and UMI sequence, retaining one representative read per unique RNA fragment.

Only the first position of the RNA fragments, meaning the 5’ end, with lengths between 28 and 30, inclusive, was kept. Additionally, the positions of the fragments were increased by one if they overlapped the second and third codon positions and had lengths of 29 and 30. For every codon in the coding sequence, the sum of the fragment counts, the peptide and nucleotide sequences in the codon, and around it were extracted for further analysis. Non-coding sequences were masked as X and N for peptides and nucleotides, respectively. Transcripts with a Reads Per Kilobase of transcript per Million mapped reads (RPKM) < 1 were discarded.

### Pseudo-bulk aggregation across replicates

Per condition, pseudo-bulk aggregates were constructed by adding the fragment counts across replicates. For each replicate, positions were indexed within a transcript, and replicate-specific values were used to align positions across replicates. We then compute the aggregated counts per transcript and position by summing the read counts across the three biological replicates for each sample. Transcripts with an average, across replicates, of Reads Per Kilobase of transcript per Million mapped reads (RPKM) < 1 were discarded.

### Ribosome stalling peak detection

To identify peaks, first, the local maximum were obtained using the function findpeaks from the pracma R package by sample. Then, for each putative peak, the start and end positions of the peaks, the local background (max read count in the +/− 25 codons around the peaks), and the global background (mean read count of the transcript excluding the peak) were computed to be used as covariates for the baseline model described below.

### Baseline model for expected read counts estimation

A baseline model for read counts using gradient-boosted regression trees is fit to quantify peak significance relative to expected local signal (gradient boosting machine (GBM) function from the gbm R package) with a Poisson response. The model predicts the expected read count for each candidate peak using the peak’s position, global background, and local background. The GBM was trained with 10-fold cross-validation and 5000 trees. Cross-validated fitted values were exponentiated to obtain the expected mean read count under the baseline model. For each candidate peak, we compute a two-sided Poisson p-value by doubling the smaller tail probability.

Multiple testing correction across all candidate peaks in a sample was perform using Benjamini-Hochberg procedure for multiple testing correction (34). A peak is called a stalling site if it has a False discovery rate (FDR) < 0.1 and log fold change (LFC) > 1.

### Gradient-boosted regression model evaluation

To evaluate how local amino acid and nucleotide sequences classify a peak as stalling and non-stalling, logistic regressions (function fastLR from the RcppNumerical R package) were fit. 5-fold cross-validated area under the receiver operating characteristic curve (AUC-ROC) curves) at different LFC thresholds.

### Stalling peaks’ sequence logos

Amino acid and nucleotide sequence logos were created for the stalling peaks (FDR < 0.1 and LFC > 1) using non-peak positions (RPKM > 0) as background. The logos are calculated as the Kullback– Leibler (KL) divergence between the stalling and background frequencies (epsilon = 1e-6) (35).

### Computational identification of G-quadruplexes

G-quadruplexes enrichment was determined around stalling and non-stalling peaks, from the same transcripts as the stalling peaks, using the G4SNVHunter R package (36). The 0 to +180 region relative to the peaks were scan for G-quadruplexes (functions G4HunterScore and G4HunterDetect, parameters: window_size = 25 and threshold = 1.2) the mean scores between groups were compared with a Wilcoxon tests (alternative = two-sided). Sampling of the non-stalling peaks was done to ensure the same number of cases are present in each group.

### Sample vitrification and tilt series data collection for Cryo-ET

Before sample vitrification, 5-nm gold fiducials conjugated with protein A (Cytodiagnostics, OD10) were added to the purified dense ribosome clusters from the pellet fraction at a volume ratio of 5:1 (samples/fiducials), yielding a final sample concentration of 575 nM. Cryo-EM grids (c-flat CF-2/2-2Cu-T) were washed in chloroform for 2 h and glow-discharged in air at 10 mA for 13 seconds before the sample was applied. Approximately 3.6 μl of sample was applied to the grid before plunge freezing into liquid ethane using a Vitrobot Mark IV (Thermo Fisher Scientific Inc.). The Vitrobot parameters for vitrification were set to a blotting time of 3 seconds and a blot force of +1. The Vitrobot chamber was maintained at 25 °C with 100% relative humidity during the process.

Tilt series were acquired on the FEMR-McGill Titan Krios electron microscope operated at 300 kV and equipped with a Gatan K3 direct electron detector and a Quantum LS imaging filter. Slit width was 30 eV. Multiple tilt series were collected simultaneously using the Parallel cryo-electron tomography (PACEtomo) script installed in SerialEM (24,25). Cryo-ET data collection parameters are described in Supplementary Table S6.

### Tomogram reconstruction and particle picking

Movies obtained for the raw tilt series were motion-corrected, and the initial CTF parameters were estimated in Warp (v2.0.0) (37). A total of 431 tilt series were further stacked in Warp before alignment. The stacks were subsequently aligned at 4x binning using in-house scripts performing fiducial tracking tilt alignment in IMOD (38). All aligned tilts were then imported into Warp for a new round of CTF parameters estimation and the tomograms were reconstructed at 15 Å/pixel for inspection and subsequent particle picking.

For template matching, an 80S ribosome map (rotated 2 state from the purified dense ribosome clusters from the pellet fraction sample) obtained from our single-particle analysis was resampled to 4.36 Å/pixel with a box size of 112 pixels, and low-pass filtered to 20 Å in cryoSPARC (26). Template matching was performed in Warp against tomograms reconstructed at 15 Å/pixel, using a rotational step of 15° with a minimum inter-particle distance of 100 Å. Finally, a global picking threshold of 2 was applied to all tomograms, yielding a total of 217,865 ribosome particles.

### 3D classification and sub-tomogram averaging

All particles identified by template matching were re-extracted in Warp at 2 x binning (3.32 Å/pixel with a box size of 148 pixels) and imported into RELION (39) for 3D classification. To curate clean particles, we performed two rounds of exhaustive 3D classification, each requesting 5 classes and 25 iterations. The regularization parameter (T) was set to 4, and a circular mask of 488 Å was used. After classification, particles corresponding to the 80S ribosomes were selected, yielding a curated dataset of 114,191 good particles.

Selected particles were pooled and subjected to a 3D auto-refinement in RELION to generate a consensus subtomogram-averaged map. To resolve the structural heterogeneity among ribosome populations, two additional layers of focused 3D classification were performed. In this step, a focused mask (4-pixel dilation, 6-pixel soft edge) was applied on the A, P and E sites of the 80S ribosome. The regularization parameter value (T) was set to 20, requesting three classes per classification layer. Particles assigned to classes representing the same features in the masked region were pooled, resulting in five distinct ribosome states. Particles from each class were independently refined using RELION’s 3D auto-refine procedures to obtain the final structures. During refinement, C1 symmetry was applied with a circular mask of 488 Å, and the map obtained from 3D classification was used as the initial model after it was low-pass filtered to 60 Å. Visualization of cryo-EM density maps was performed in UCSF Chimera and Chimera X (28,29). Final subtomogram averages were mapped back to the original tomograms using ArtiaX (40), which was also used for the downstream structural and statistical analyses.

### 3D classification and multibody refinement of disome structures linked by rRNA expansion segments

To determine the relative orientation of ribosomes from different polysomes associated through rRNA expansion segments, and to obtain cryo-EM structures of the corresponding disomes, we selected all disomes composed of two ribosomes in the rotated-2 state from tomograms with a 4 Å/pixel sampling. Disomes were extracted in Warp using a box size of 246 pixels and imported into RELION (39) for refinement.

We first performed a consensus refinement using the subtomogram-averaged cryo-EM map of the rotated-2 ribosome, low-pass filtered to 20 Å, as the initial reference. Refinement was carried out with a 984 Å circular mask, an initial angular sampling of 7.5°, an offset range of 5 pixels, an offset step of 1 pixel, and local angular searches from autosampling at 1.8°. This refinement aligned the left ribosome within the disome well; however, the right ribosome appeared as poorly defined, amorphous density, indicating variability in its orientation relative to the left ribosome.

To resolve this conformational heterogeneity, we performed focused 3D classification on the density corresponding to the right ribosome. A mask was applied around this region, and the same low-pass-filtered rotated-2 ribosome map was used as the initial reference. Classification was performed into five classes using a regularization parameter of T=20T=20T=20, an angular sampling interval of 7.5°, an offset range of 5 pixels, and an offset step of 1 pixel. Classes in which the density resembled an 80S ribosome were selected, combined, and subjected to a second round of focused 3D classification using the same parameters, but with three requested classes.

For each of the three resulting disome classes, the right ribosome density was locally refined using an initial angular sampling and local search interval of 1.8°. We then repeated local refinement for the left ribosome in each class, yielding the final cryo-EM maps of three distinct disome arrangements.

The multibody refinement (41) of these disomes was performed using the orientation parameters of the consensus refinement as the starting point, and each of the ribosomes in the pair was defined as a ‘body’. The multibody refinement was run using an initial angular sampling of 3.7 degrees, an initial offset range of 3 pixels, an initial offset step of 0.75 and requested 5 eigenvector movies.

## RESULTS

### Half of the ribosomes included in the dense ribosome clusters are stalled in elongation

We purified ribosomal clusters from homogenized P5 (5-day-old) rat brains using sucrose gradient ultracentrifugation (20). At this developmental stage, over 90% of brain cells are neurons, minimizing contamination from glial ribosomes (42). As shown in previous publications (20), this protocol effectively separates the dense ribosomal clusters, often associated with neuronal RNA granules (43), which are deposited in the pellet, from ribosomes included in other cellular ribosome fractions, such as polysomes (migrating in fractions 5/6) or monosomes (migrating in fractions 2/3) (Figure 1A). Negative staining electron microscopy confirmed the enrichment of ribosome clusters (Supplementary Figure S1).

**Figure 1.**
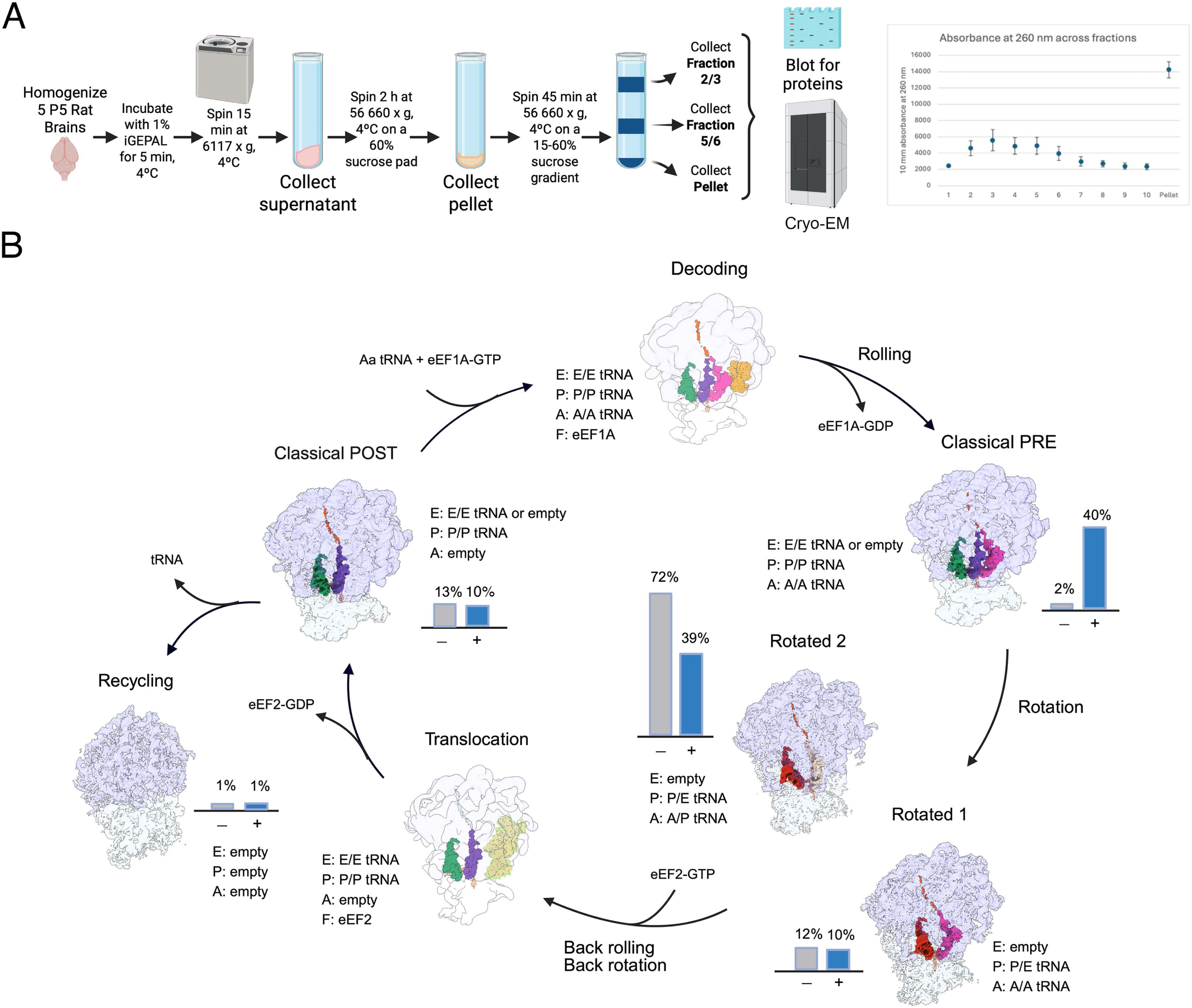
Elongation intermediates in purified neuronal RNA granules. (A) Schematic depicting the purification steps for the dense ribosome clusters in the pellet fraction, and in fractions 5/6, and 2/3 from P5 rat brains. These fractions were analyzed using western blotting and cryo-EM in this study. UV absorbance of the sedimentation through the sucrose gradient is shown in the right panel). (B) The cryo-EM maps of elongation intermediates obtained through image classification from purified dense ribosome clusters from the pellet fraction, are positioned within the canonical elongation cycle. Densities representing the 40S and 60S subunits are displayed as transparent to highlight the tRNA molecules within the ribosome for each elongation state. Each elongation intermediate is characterized by the location of the tRNA in the A, P, and E sites. The bar graphs beside each map indicate the proportion of particles in that elongation state before (grey) and after (dark blue) the sample was treated with cycloheximide. Main movements of the ribosomal particles during the cycle (rotation, back rotation, rolling, and back rolling) are marked along the cycle. Elongation states for which particles were not isolated from the dataset are shown as faded structures. The ‘decoding state’ was created using the EMDB-2624 map and the PDB ID 4cxh molecular model. The ‘translocation state’ was created using the EMDB-38666 map and the PDB ID 8yld molecular model.

Recent cryo-EM analysis of dense ribosome clusters (20) revealed that ∼85% of the ribosomes included in the clusters were stalled at elongation and contained a nascent polypeptide in the exit tunnel. A caveat in this initial structural analysis was that the buffers used in these purifications lacked spermine and spermidine. These conditions can cause dissociation of ribosomal subunits and tRNAs (44), resulting in a skewed description of the ribosome populations within the dense ribosome clusters and the state of the translation cycle in which they are found (45).

To obtain a more comprehensive and accurate description of ribosome populations in neuronal ribosome clusters, we used a modified purification protocol. The protocol used buffers containing 0.04 mM Spermine, and 0.5 mM Spermidine, which help preserve the structural integrity of translation intermediates (46), and avoided the high-salt step used earlier to facilitate cleavage of clusters into monosomes (20). We then examined RNase I-treated purified monosomes by cryo-EM (Supplementary Table S1) and processed the images with both global and focused 3D classification to identify all elongation intermediates. The cryo-EM maps for these elongation intermediates were refined to resolutions ranging from 2.4 to 4.7 Å (Supplementary Figure S2 & Supplementary Table S1). We found that ribosome clusters from the pellet contain ribosomes in multiple elongation states (Figure 1B). The majority (∼84%) were in the rotated states, consistent with previous research (20,47). Specifically, 12% of ribosomes were in the rotated 1 state, while 72% adopted the rotated 2 state. Ribosomes in the classical POST state comprised 13%, and 2% were in the classical PRE state. Approximately 1% of the ribosomes lacked any visible tRNA densities, representing recycling intermediates or dormant ribosomes.

Consistent with the protective effect of spermine and spermidine in the homogenization and purification buffers, we observed less than 1% free 60S subunits (not shown in Figure 1B), compared with 13% in our previous studies (48). In addition, these buffers also preserved the tRNA in the E site (49), which was present in all elongation states, but was not observed previously in the classical PRE or POST states (20,48).

The elongation states observed in this study showed almost a complete absence of density for the P stalk in the 60S subunit. The density for the uL11 (Rpl12) was highly fragmented, and there was no density for uL10 (Rplp0). These are the two ribosomal proteins that form the base of the P stalk and recruit the eEF2 elongation factor. Densities representing these two proteins were also absent in the cryo-EM maps of the elongation intermediates observed in our previous studies (20,48). Therefore, these new structures allow us to conclude that the high flexibility of the P stalk in the elongation intermediates included in the dense ribosome clusters was not a consequence of the absence of polyamines in the buffers. These new results indicate that prevention of eEF2 may be a part of the mechanism to halt translation on these ribosomes.

To determine the fraction of these elongation intermediates that are stalled prior to homogenization, we repeated the purification and cryo-EM analysis (Supplementary Table S2) of the dense ribosome clusters in the pellet fraction, this time including 0.1 mg/ml cycloheximide (CHX), an elongation inhibitor, in the homogenization and purification buffer. CHX binds to the E site on the large ribosomal subunit and prevents tRNA accommodation at this site. In elongating ribosomes, CHX occupancy at the E site arrests all translating ribosomes in the classical PRE state, with the nascent peptide transferred to the A-site and preventing the transition to rotated configurations (50). Consequently, ribosomes in the classical PRE state containing a CHX in the E site after CHX treatment originate from actively elongating ribosomes, whereas ribosomes in other states are, by definition, pre-stalled prior to the treatment.

We found that the percentage of ribosomes in the classical-PRE state increased from 2% to 40% (Figure 1B & Supplementary Figure S3A). The CHX molecule was also visible in the E site of the large subunit, confirming its inhibitory activity (Supplementary Figure S3A). However, for the rotated 2 state ribosomes, the percentage decreased to 39%, and for the rotated 1 state ribosomes, it also decreased to 10%. A molecule of CHX was not visible in the E site of the large subunit of the rotated 1 and rotated 2 ribosomes (Supplementary Figure S3A). About 10% of ribosomes were in the classical-POST state, containing a P/P tRNA but lacking an E/E tRNA. Instead, a density for CHX was visible in the E-site (Supplementary Figure S3A). This percentage was largely unchanged compared to the untreated sample. Approximately 1% of the ribosomes also lacked any visible tRNA densities, representing recycling intermediates or dormant ribosomes. The cryo-EM maps for these elongation intermediates were refined to resolutions ranging from 2.8 to 4.9 Å (Supplementary Figure S4 & Supplementary Table S2).

Overall, these results indicate that about 49% of the ribosomes in dense ribosome clusters are stalled during elongation. Importantly, the addition of cycloheximide confirmed that stalling was not solely due to runoff, as this compound reduces the percentage of hybrid-state ribosomes but still leaves half the ribosomes in the rotated state.

### Fractions equivalent to the polysome and monosome fractions also include ribosome clusters containing stalled ribosomes

In our purification protocol (Figure 1A), fractions 5/6 of the final sucrose gradient sedimentation contain ribosome clusters comparable in size to polysomes in other cells (51,52). However, images of the ribosomes contained in fractions 5/6 obtained by negative staining electron microscopy show ribosome clusters similar to those found in the pellet fraction after resuspension (Supplementary Figure S1B). These clusters were markedly different in morphology from polysomes containing actively translating ribosomes in other cells from the same animals, such as hepatocytes. Polysomes from hepatocytes were mainly linear arrays of ribosomes along a single mRNA molecule, often adopting a circular shape (Supplementary Figure S1D). We also analyzed fractions 2/3 from our gradients, which are equivalent to the monosome/disome fraction obtained when purifying ribosomes from non-neuronal cells. In this case, images revealed that in addition to ribosome clusters, there were also abundant monosomes and disomes (Supplementary Figure S1C).

In most cells, polysomes represent the fraction of ribosomes actively translating proteins (53). To test whether the ribosomes in fractions 5/6 of our gradient were also actively translating, we subjected this sample to the same cryo-EM analysis that was performed with the pellet fractions (Supplementary Table S3). In the absence of CHX, the distribution of elongation intermediates was very similar to that found in the pellet fraction, suggesting also a high proportion of stalled translation intermediates. The rotated 1 (13%) and rotated 2 (72%) states were the most abundant, followed by the ribosomes in the classical POST state (11%) and classical PRE state (2%). Finally, 2% of the ribosomes were recycling intermediates lacking visible tRNA densities (Figure 2). The cryo-EM maps for these elongation intermediates were refined to resolutions ranging from 2.4 to 4.5 Å (Supplementary Figure S5 & Supplementary Table S3).

**Figure 2.**
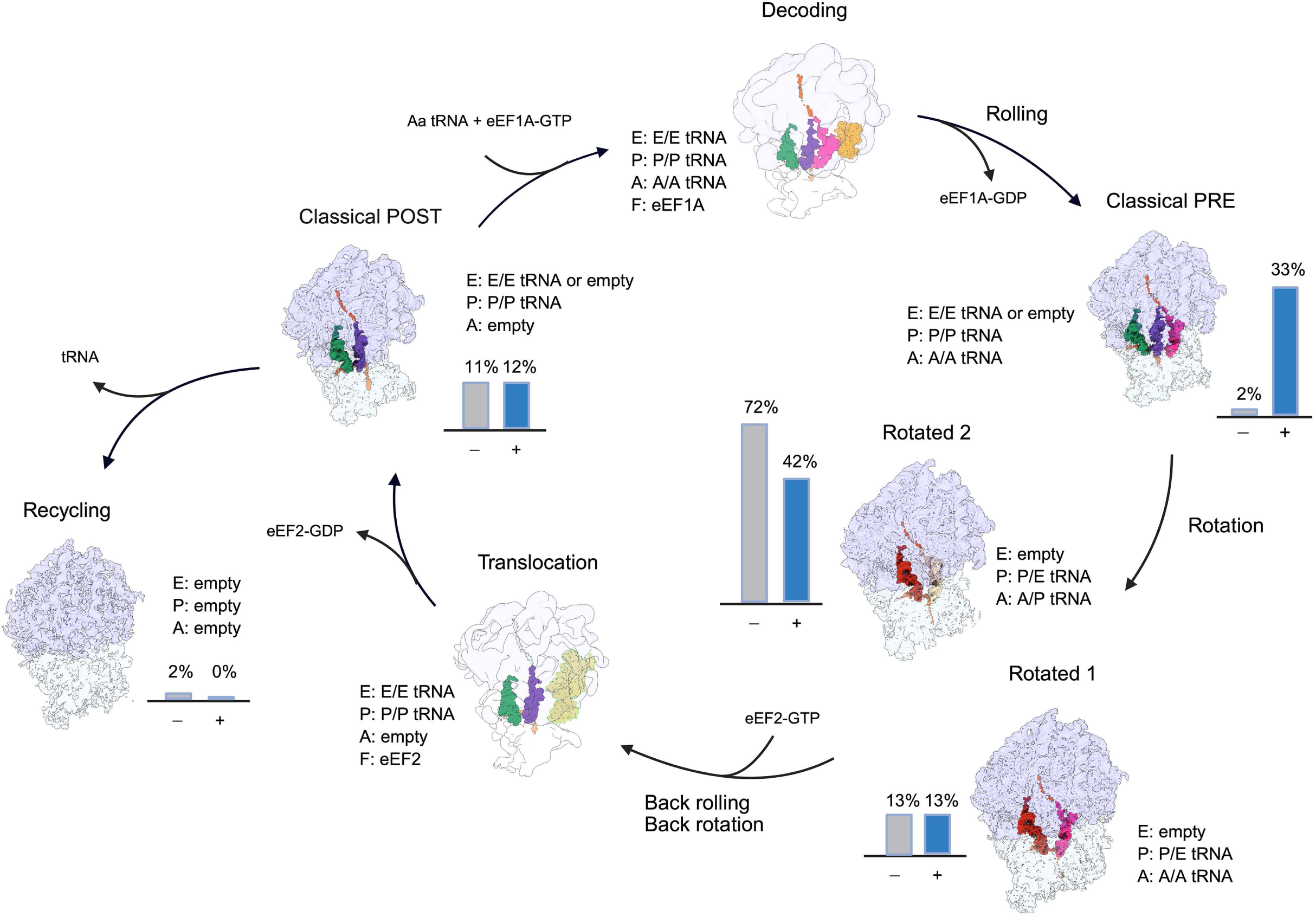
Elongation intermediates in fractions 5/6. (A) Cryo-EM maps of elongation intermediates obtained through image classification from fractions 5/6, positioned within the canonical elongation cycle. The layout of this panel and labels are as in Figure 1, panel (B). The bar graph beside each map indicates the proportion of particles in that elongation state before (grey) and after (dark blue) the sample was treated with cycloheximide.

In contrast, when fractions 2/3 were analyzed (Supplementary Tables S5), we found that the rotated 2 and rotated1 states reached only 38% and 9%, respectively (Figure 3). Classical POST state intermediates accounted for 16%. However, most of the remaining ribosomes (37%) were in recycling (containing only an E/E tRNA and an absent nascent polypeptide), suggesting that these ribosomes were active but had undergone run-off during brain homogenization and had completed translation of the mRNA. These ribosomes were likely formed from non-stalled ribosomes translating, either as a monosome or a disome, and thus were able to run off the mRNA and enter the recycling state. This is consistent with the higher proportion of monosomes and disomes in fractions 2/3 (Supplementary Figure S1C). Non-stalled ribosomes that were part of longer polysomes or ribosome clusters during brain homogenization could not evolve to a recycling state because they collided with the downstream stalled ribosomes within the same mRNA, remained associated with the mRNA, and became stalled into a rotated state, contributing to increasing the percentage of rotated-1 and rotated-2 stages that migrated into fractions 5/6 and pellet. The cryo-EM maps for the elongation intermediates found in fractions 2/3 were refined to resolutions ranging from 2.6 to 3.3 Å (Supplementary Figure S7 & Supplementary Table S5).

**Figure 3.**
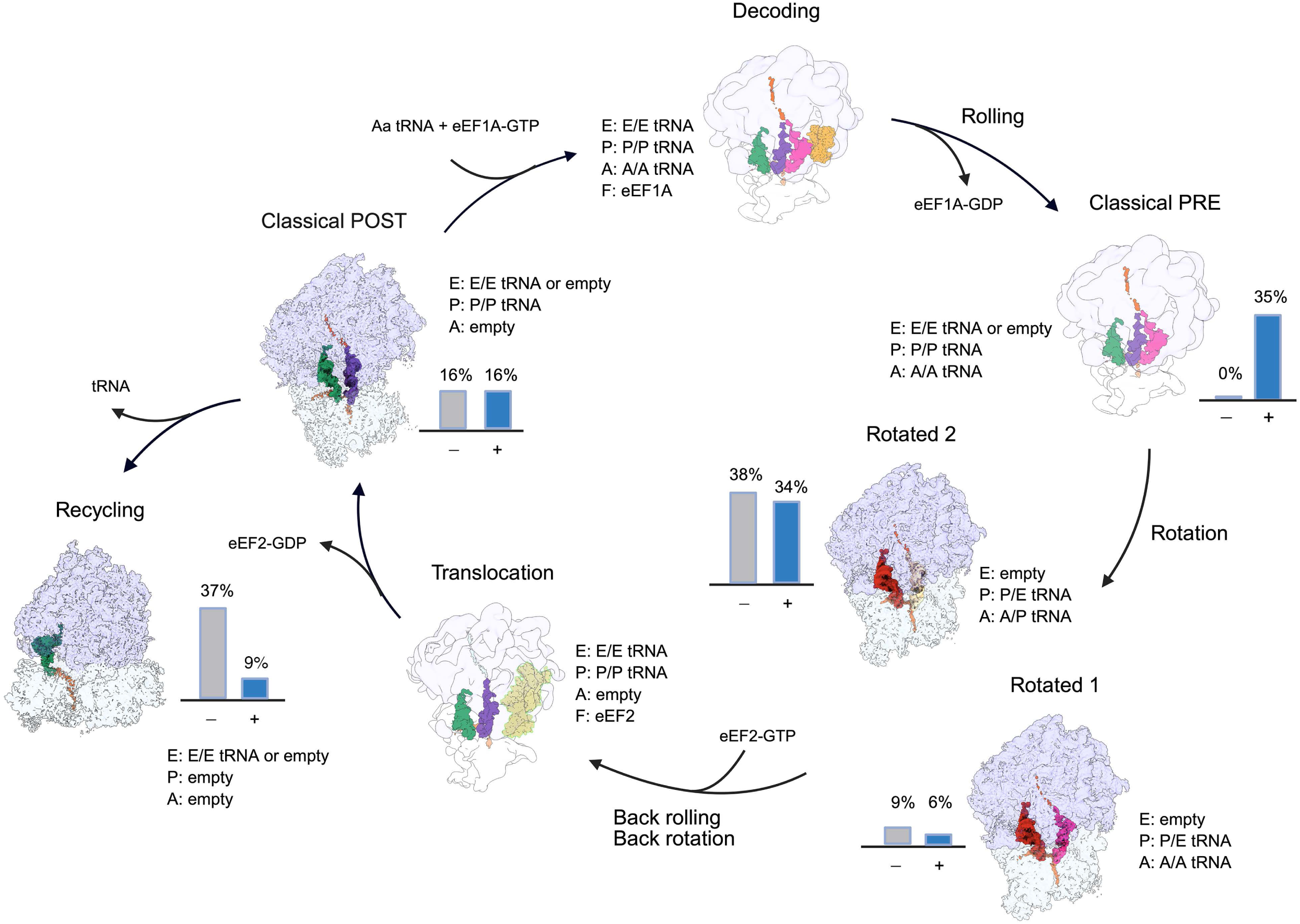
Elongation intermediates in fractions 2/3. (A) Cryo-EM structures of elongation intermediates, obtained through image classification from fractions 2/3 positioned within the canonical elongation cycle. The layout of this panel and labels are as in Figure 1, panel (B). The bar graph beside each map indicates the proportion of particles in that elongation state before (grey) and after (dark blue) the sample was treated with cycloheximide.

To more accurately quantify the percentage of stalled ribosomes in fractions 5/6 and 2/3, we repeated their analysis after CHX treatment. In fractions 5/6, the addition of CHX showed effects similar to the pellet fraction (Figure 1B). The percentage of the rotated 2 state decreased by approximately half to 42%, and the percentage of the rotated 1 state remained at 13% (Figure 2), indicating that about half of the ribosomes in the fractions 5/6 were also stalled. In addition, the percentage of ribosomes in the classical-PRE state increased from 2% to 33%, driven by the active translation intermediates that stall upon CHX binding in this state, preventing them from transitioning into the rotated state. The percentage of classical POST state ribosomes did not change significantly, changing to 12%. Similar to the pellet fraction, the molecule of CHX was visible in the E site of the classical PRE and classical POST states, but not in the rotated states (Supplementary Figure S3B). The cryo-EM maps for these elongation intermediates were refined to resolutions ranging from 2.5 to 2.8 Å (Supplementary Figure S6 & Supplementary Table S4).

In fractions 2/3, the CHX treatment only slightly dropped the proportion of the rotated 1 and rotated 2 states to 6% to 34% ribosomes, respectively, indicating that also in this fraction, about half of the ribosomes were also stalled. However, CHX blocked actively translating ribosomes, likely in monosome or disome form, from running off into the recycling state, reducing it to 9%. It captured this population instead into the classical PRE state, which increased from 0% to 35%. The proportion of intermediates in the classical POST did not change and remained in 16%. As in the other CHX-treated samples, the molecule of CHX was visible in the E site of the classical PRE and classical POST states, but not in the rotated or recycling (Supplementary Figure S2C) states. The cryo-EM maps for these elongation intermediates were refined to resolutions ranging from 2.7 to 3.4 Å (Supplementary Figure S8 & Supplementary Table S6).

Overall, these results indicate that the ribosome populations in fractions 5/6 and 2/3 are similar to those in the pellet fraction, and that, in both samples, almost half of the ribosomes were stalled in elongation. Given the different sedimentation velocity of the dense ribosome clusters in the pellet fraction compared to the intermediates in fractions 5/6 and 2/3, these data suggest that ribosomes in these fractions may represent the same ribosome population but differ in the organization of the ribosome clusters.

### Distribution of RNA-binding proteins between the translating and stalled fractions

We previously examined the distribution of several RBPs in the neuronal pellet and fractions 5/6. However, we have not examined the distribution of RBPs in the lighter fractions 2/3 (Figure 1A). As discussed, the buffers used in our previous study (20) lacked spermine and spermidine, which may have caused dissociation of RBP from ribosome clusters.

To determine if there are major differences in the fractionation of RBPs using the buffers containing spermine and spermidine, we repeated this analysis (Figure 4A). Notably, hnRNPA2/B1, an RBP strongly linked to RNA transport through binding to 3’UTR of selected messages (54), was strongly enriched in fractions 2/3 compared to fractions 5/6 and the pellet. In contrast, some RBPs previously implicated in the transport of stalled ribosomes, such as Staufen 2 and FMRP, are highly enriched in fractions 5/6 and the pellet compared to fractions 2/3. Many other RBPs, such as G3BP1, PurA, and UPF1, were found distributed throughout the fractions. The level of eEF2 was low, particularly in fractions 5/6 and the pellet, consistent with the fragmented cryo-EM density observed for the P stalk and with the role of this structural motif in recruiting the elongation factor. Consistent with the cryo-EM analysis suggesting that intermediates in fractions 5/6 and pellet are similar but exhibit different degrees of compaction, we found that using the new buffer with polyamines, there were only small differences between the pellet and fractions 5/6 after normalization to S6 compared to previous studies (20,55). For the RBPs, we determined whether there were significant differences in percentages across the fractions. hnRNPA2/B1 had a significantly higher percentage in fractions 2/3 compared to most of the RBPs, while Staufen 2 had a significantly higher percentage in the pellet compared to some RBPs (Figure 4B).

**Figure 4.**
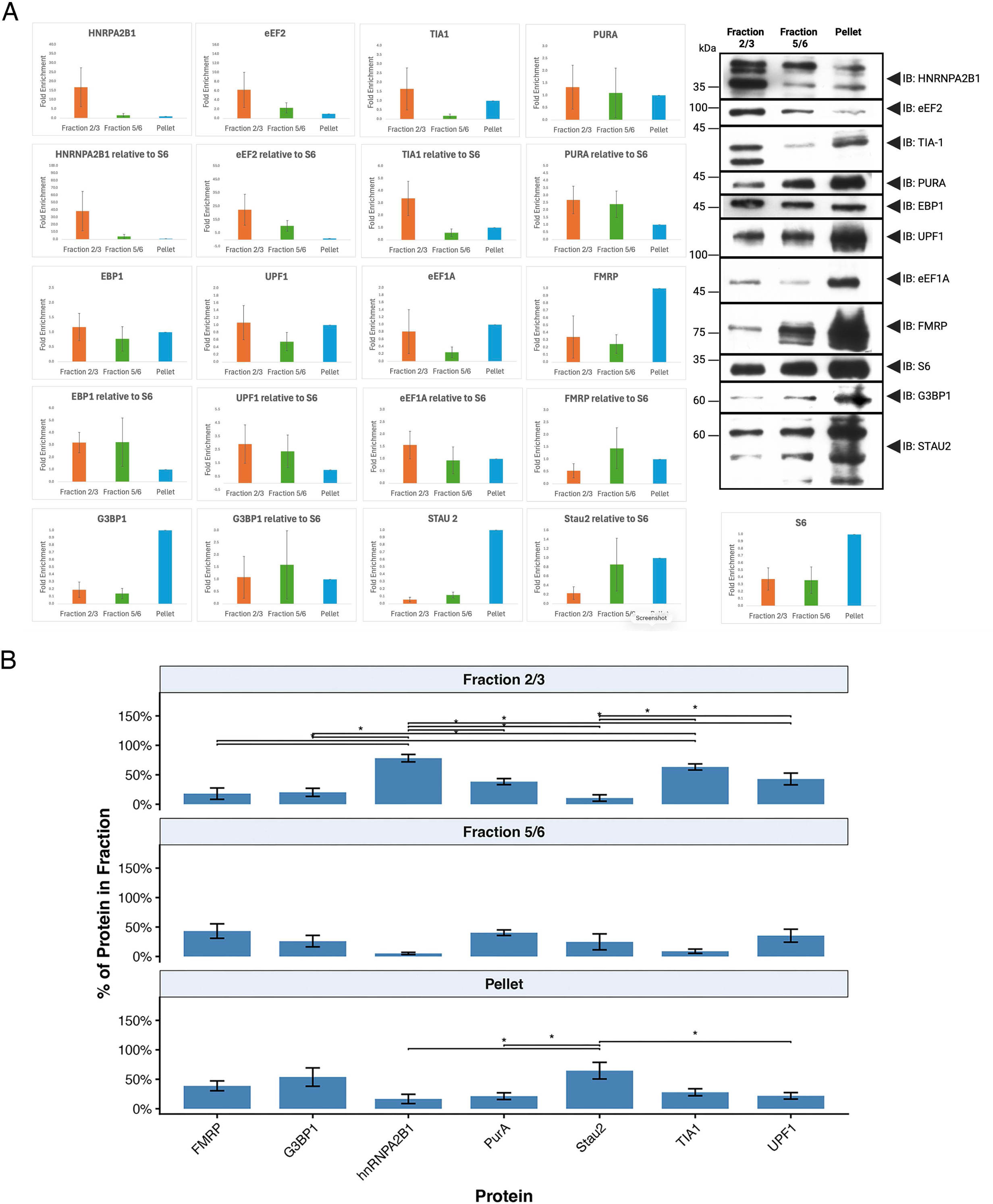
Immunoblot analysis of fractions 2/3, 5/6, and pellet. We stained for hnRNPA2B1, eEF2, TIA1, PURA, EBP1, UPF1, eEF1A, FMRP, S6, G3BP1, STAU2. One representative blot for each protein is shown in the right panel. The bar graphs show the quantification results of the Western blots. The fold enrichment of each fraction compared with the pellet is shown for each protein, in order of enrichment of fractions 2/3: hnRNPA2B1 (N=4), eEF2 (N=3), TIA1 (N=5), PURA (N=5), EBP1 (N=5), UPF1 (N=5), eEF1A (N=5), FMRP (N=4), S6 (N=5), G3BP1 (N=3), STAU2 (N=3). Errors are S.E.M. B) To compare the percentage of RBPs in each fraction, the data was recalculated by dividing the normalized levels of each protein by the total normalized protein of that protein in the three fractions for each replicate. Errors are S.E.M. ANOVAs were done for each fraction. Fraction 2/3 F(32,6)=12.1, p< 0.1E-05; Fraction 5/6 (F32,6)=3.0. p<0.02; Pellet (F(32,6) <0.01. Tukey Post-Hoc tests were done for each fraction and significantly different groups are shown in the figure (*, p<0.01).

### Ribosomes in the clusters are stalled at specific sequences

It is still unknown what information in the mRNA determines where stalling occurs. To analyze this aspect of the mechanism, we analyzed ribosome-protected fragment (RPF) data from our previous work (55). RPFs are typically 28-30 base pairs (Figure 5A). We counted the number of RPFs that started on each codon and used a gradient-based regression learning model to determine the increase in RPFs at specific sites over expected RPFs to identify stalling sites (See Methods). We then used logistical regression to determine whether sequences in the RPFs and the surrounding sequence could predict the stalling sites. We found that the best-performing model (highest area under the curve) combined information from amino acids at the entrance to the peptide exit tunnel and from nucleotides throughout the sequence. Similar results were obtained for RPFs from the pellet (Figure 5B, top left panel) and fractions 5/6 (Figure 5B, top right panel). Finally, by comparing the logistic regression results with the expected background based on amino acid frequencies and codon usage, we generated iceLOGOs that identified nucleotides and amino acids enriched at stalling sites in the pellet and fractions 5/6 (Figure 5B, middle panel).

**Figure 5.**
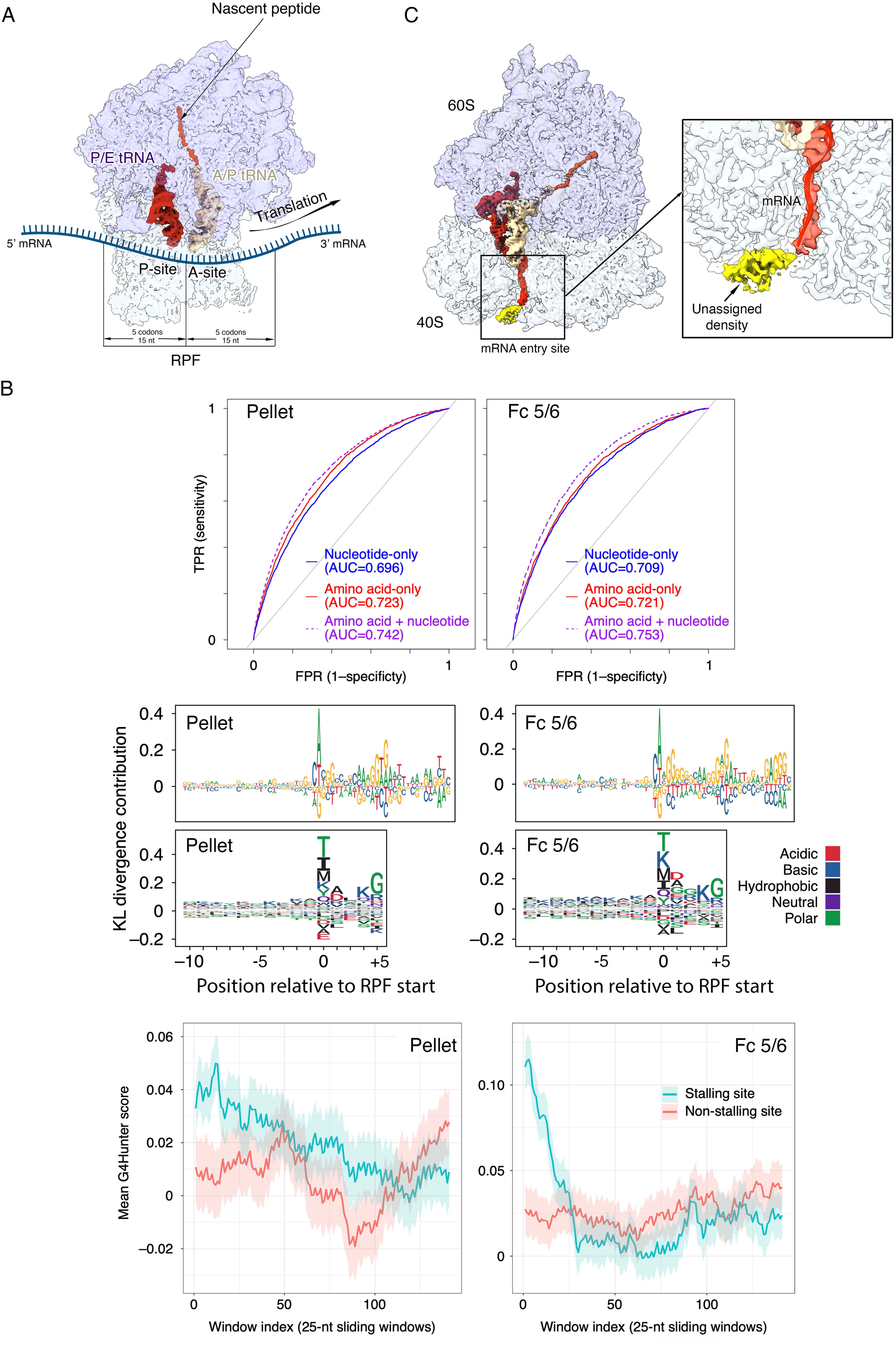
Figure showing sequence analysis and extradensity in the mRNA entrance. (A) Cartoon representing the ribosome in the rotated 2 state while translating a mRNA molecule. The A/P and P/E tRNAs and the nascent peptide in the exit channel are shown. The diagram also shows the ribosome protected fragment (RPF) of the mRNA molecule and the distances in nucleotides (nt) and codons from the beginning of the RPF to the location of the P site and end of the RPF. (B) Top panels: Receiver Operating Characteristic (ROC) curves demonstrating the performance of logistic regression models trained on nucleotide sequences (blue), amino acid sequences (red), or both (purple) to separate stall sites from non-stall (background) positions in the ribosomes from the pellet fraction (left panel) and fractions 5/6 (right panels). TPR: true positive rate. FPR: false positive rate. Metrics are based on 5-fold cross-validation. Middle panels: Nucleotide (top) and amino acid (bottom) iceLOGOS summarizing sequence patterns at and around stall sites in the ribosomes from the pellet fraction (left panel) and fractions 5/6 (right panels). Letter heights correspond to point-wise contribution to KL divergence relative to background non-stall sites; positive values indicate enrichment in stall sites relative to background, and negative values indicate depletion. Positions are relative to the start of the RPF and are expressed in codons. Bottom panels: Moving-window average propensity to form G-quadruplex (G4) by stall (cyan) and non-stall (light red) sites in the ribosomes from the pellet fraction (left panel) and fractions 5/6 (right panels). Positions, in nucleotides (nt), are relative to the RPF start site, representing the beginning of each 25-nt window whose mean score is shown. Shaded areas represent the standard error of the mean. A Wilcox test was performed to compare the stalling and non-stalling groups (M_WT_fractions 5/6 p-value = 4.7e-5, and M_WT_pellet p-value = 0.15. (C) Cryo-EM map of the rotated 2 elongation intermediates from purified dense ribosome clusters from the pellet fraction. The boxed area in the left panel indicates the mRNA entry site, and the right panel shows a zoomed-in view of this area, displaying the unassigned density observed at the mRNA entry point into the ribosome, caused by this mRNA fragment forming a G-quadruplex structure or another alternative secondary structure.

The largest amount of information is close to the RNA exit site where an A nucleotide is favored in the iceLOGOs for the pellet (Figure 5B, middle left panel) and fractions 5/6 (Figure 5B, middle right panel). This appears to partially drive the amino acid in this position as the codons encoding the four most likely amino acids here (T, I, M and K) all start with an A nucleotide. We do not detect any feature at the RNA exit site or in the nascent peptide exit channel in the cryo-EM structures that could explain this preference. In the nucleotide iceLOGOs we also noted an enrichment for Gs in the nucleotides following the expected codon in the A site. Using computational methods to predict G quadruplexes (36), we showed that stalling sites had a higher propensity to form G quadruplexes than non-stalling sites (Figure 5B, bottom panel). Interestingly, we observed in the cryo-EM maps of most elongation intermediates from the pellet and fractions 5/6 an unassigned density near the mRNA entry site (Figure 5C). This region in our current cryo-EM map lacks sufficient local resolution to unambiguously assign this density to a specific RNA structure. However, it could represent an RNA G-quadruplex, which forms in guanine-rich RNAs. Thus, the places in the mRNA where stalling occurs appear to be driven by a combination of factors, including a secondary structure near the mRNA entry site, specific nucleotides at the mRNA exit site, and amino acids at the beginning of the peptide tunnel.

### Ribosome clusters in neuronal RNA granules contain multiple polysomes

In the cryo-EM analysis described in previous sections, the dense ribosome clusters in the pellet fraction were treated with nuclease before imaging to separate the ribosomes within the clusters into individual monosomes that can be studied by single-particle approaches. However, the higher-order organization of the ribosomes on the clusters was lost with this approach. To study this aspect of the dense ribosome clusters in the pellet, we collected cryo-electron tomography data (Supplementary Table S7) on these clusters without nuclease treatment and performed sub-tomogram averaging of the 80S ribosome particles (Supplementary Table S7 and Supplementary Figures S9 and S10). Overall, we found that the percentages of the multiple elongation intermediates within the clusters without nuclease treatment (Supplementary Figure S9) closely resembled those observed in the nuclease-treated dense ribosome clusters analyzed by single-particle approaches (Figure 1B). Most of the ribosomes were in the rotated 2 state (75%). The percentage of the classical POST (10%) was similar, but the proportion of classical PRE was slightly higher (11%), and the proportion of the rotated 1 state was slightly lower (4%). Overall, these results validated that the nuclease treatment did not affect the proportion of elongation intermediates in the dense ribosome clusters from the pellet fraction.

Next, we mapped the subtomogram averages (Figure 6B) of the elongation intermediates back into the 3D tomograms to examine their distribution within the ribosome clusters (Figure 6A). Consistent with the negative-staining EM results (Supplementary Figure S1A), we found that most ribosomes were part of compact clusters (Figure 6A), distinct from canonical polysomes (Supplementary Figure S1D), which typically adopt helical, zigzag, or circular conformations (51,56). These clusters had no uniform morphology and varied widely in size and shape. There were also a few monosomes and disomes outside the main clusters–likely detached during plunge-freezing vitrification or resuspension of the pellet.

**Figure 6.**
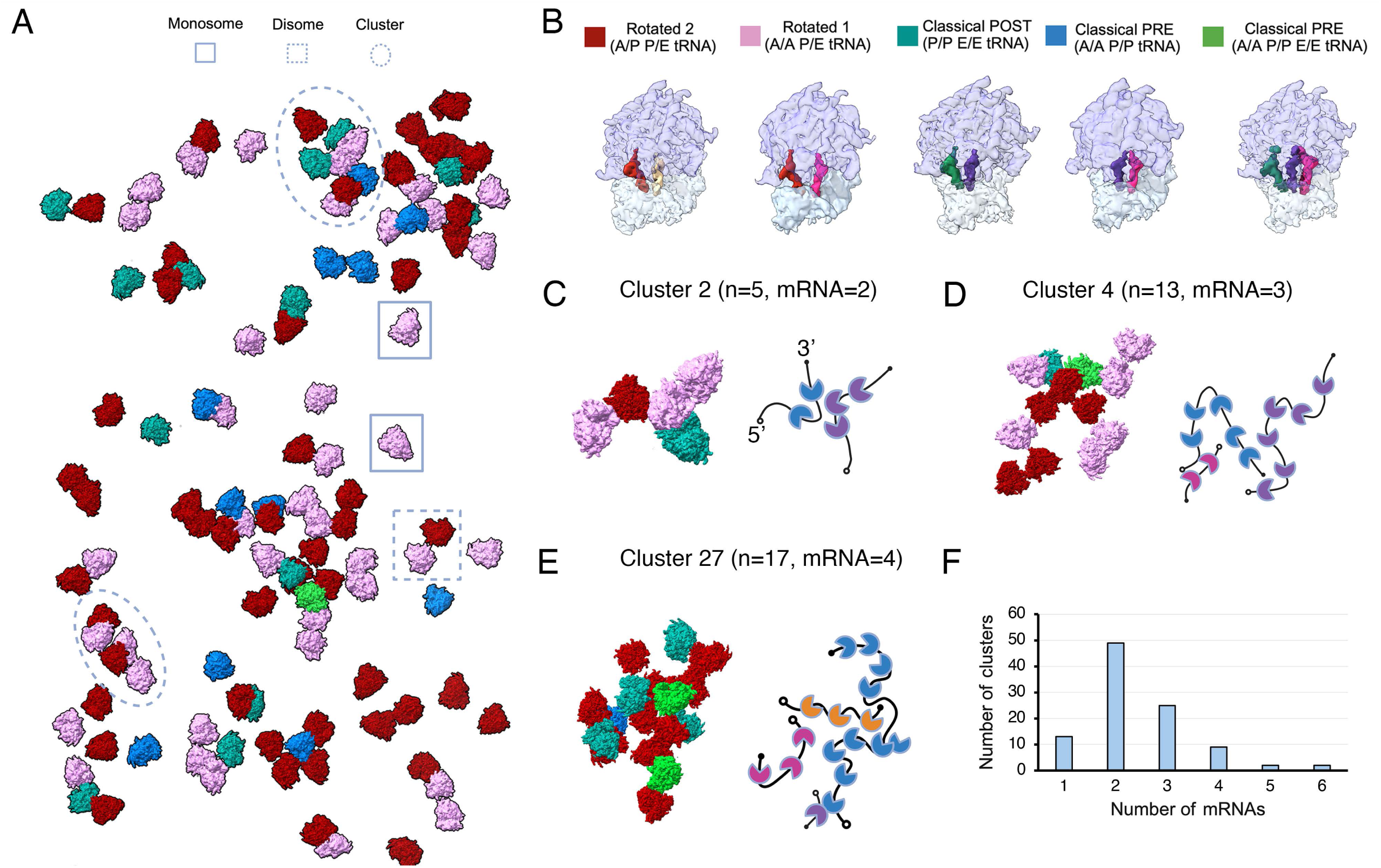
Analysis of the dense ribosome clusters from the pellet fraction by cryo-electron tomography. (A) Overview of ribosomes identified in a representative tomogram of purified dense ribosome clusters from the pellet fraction. Individual ribosomes in the tomogram are visualized by mapping back the subtomogram-averaged structure corresponding to their assigned elongation-state class, using the color scheme shown in (B). Each ribosome in the cluster is represented by a subtomogram average obtained by averaging multiple individual ribosomes in the same elongation state across all tomograms in the dataset. Examples of what constitute a monosome, disome, and cluster of ribosomes are indicated. (B) Close-up view of the subtomogram averages obtained for each elongation state from all the tomograms in the dataset. Densities representing the 40S and 60S subunits are displayed as transparent to highlight the tRNA molecules within the ribosome for each elongation state. Each elongation intermediate is characterized by the location of the tRNA in the A, P, and E sites. The color assigned to each elongation state and the positions of the tRNAs inside each ribosome is indicated at the top of each intermediate. (C-E) Analysis of the number and path of mRNA molecules in clusters 2, 4 and 27. The number of ribosomes (n), and mRNAs in each cluster is indicated. The schematic next to each cluster described the path of the mRNA molecules in each cluster. The 5’ and 3’ end of the mRNA is indicated. (F) Bar diagram indicating the number of mRNA molecules found in the 100 selected clusters.

We randomly selected 100 ribosome clusters from 38 tomograms (Supplementary Figure S10A) and analyzed their organization. The selected clusters contained between 3 and 15 ribosomes, and these were in all possible elongation intermediate states (Supplementary Figure S10A and S10B). We first determined whether the clusters contained one or multiple mRNAs. In canonical polysomes, multiple ribosomes translate the same mRNA from the 5’ to the 3’ end. The ribosome nearer to the 3’ end is regarded as the leading ribosome, while the one closer to the 5’ end is the trailing ribosome. The approximate path of the mRNA within these ribosome clusters can be estimated by the positions of the mRNA entry and exit sites on the ribosomes inside a cluster (52,57). When the mRNA exit site of one ribosome is aligned near (∼85 Å) the mRNA entry site of an adjacent ribosome, the former is typically the leading ribosome, and the latter the trailing one. Using this approach, we traced the putative mRNA paths and identified ribosomes reading the same mRNA. For example, cluster 2 contained five ribosomes (Figure 6C). Two of them followed one mRNA path, whereas the remaining three aligned with a separate mRNA. Cluster 4 (Figure 6D) contained up to 6 ribosomes along a single mRNA, and cluster 27 up to 10 (Figure 6E). Extending this analysis to the 100 selected clusters showed that 87% of the clusters contained multiple mRNA molecules (Figure 6F). This analysis allowed us to conclude that ribosomes included in the dense ribosome clusters in the pellet fraction are not a single stalled polysome, but rather a heterogeneous condensate composed of multiple polysomes.

### Ribosomes in different polysomes within the dense ribosomal clusters interact through rRNA expansion segments

In 85% of the clusters containing multiple polysomes, some ribosomes reading one of the putative mRNA molecules were positioned close enough to interact with other ribosomes reading a different mRNA (Figure 7A, left and middle panel). Examining these ribosomes across the selected clusters, we found that these ribosomes formed contacts via two structural motifs on the solvent-exposed side of the large subunit, specifically in the region beneath the L1 stalk. The corresponding cryo-EM densities resembled an A-form RNA helix. Docking a rat 80S model (47) into the density maps identified the interacting protrusions as rRNA expansion segments ES31L helix b (ES31Lb) and ES20L helix a (ES20La). In every interface observed, ES31Lb from one ribosome approached ES31Lb from the opposing ribosome, and ES20La similarly paired with its counterpart in the second ribosome (Figure 7A, right panel). Consistent with the rotated 2 state being the predominant elongation intermediate in these ribosome clusters, approximately 52% of the observed ribosome pairs were formed between two ribosomes, both in the rotated 2 state, and in 38%, at least one of the interacting ribosomes was in the rotated 2 state. Although rotated 2 ribosomes were the most frequently observed in these ribosome pairs, all other elongation-state combinations were also observed across the dataset, albeit at substantially lower frequencies. This indicates that formation of this interface is strongly enriched in the rotated 2 state but is not strictly restricted to a specific elongation-state pairing.

**Figure 7.**
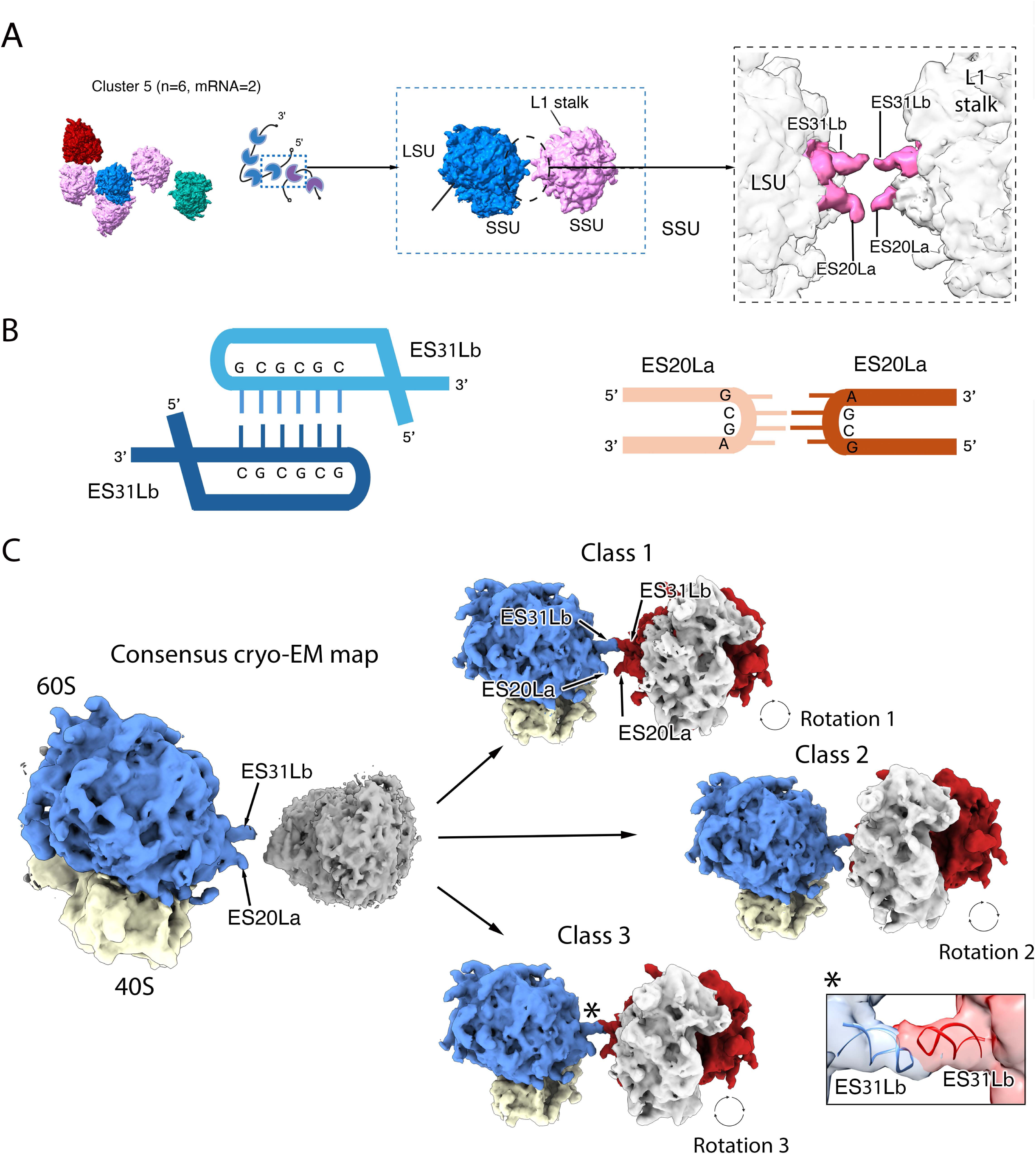
Interactions between ribosomes from different polysomes in the dense ribosomal clusters from the pellet fraction. (A) Subtomogram averages of the elongation intermediates present in cluster 5 are shown in the top panel, alongside a schematic illustrating the trajectories of the two mRNA molecules within the cluster. The dashed box highlights two ribosomes translating different mRNA molecules that contact one another through kissing-loop interactions formed by the rRNA expansion segments ES31Lb and ES20La. (B) Schematic diagrams of the kissing-loop interactions mediated by pairs of ES31Lb and ES20La expansion segments, respectively. Each diagram depicts the unpaired loop nucleotides within the corresponding rRNA expansion segment and illustrates how complementary loop–loop contacts can form kissing-loop interactions. (C) Consensus cryo-EM map obtained by refining all ribosome pairs in which both ribosomes adopt the rotated 2 state. The three disome classes identified by image classification are shown on the right. Across these classes, the ribosome on the right adopts distinct orientations relative to the ribosome on the left, revealing conformational variability within the interaction interface. A zoomed-in view of the ES31Lb: ES31Lb interaction in class 3 (indicated with an asterisk) is shown with a molecular model of these rRNA expansion segments obtained from PDB 6QZP.

Rat ES31Lb and ES20La adopt hairpin conformations. Each hairpin terminates in a loop of unpaired nucleotides: four in ES20La and six in ES31Lb. In ES20La, the loop sequence is GCGA (nucleotides 2862–2865, human numbering), whereas in ES31Lb it is GCGCGC (nucleotides 4139–4144). This structural arrangement enables the terminal loops of these expansion segments to mediate inter-ribosomal kissing-loop interactions (58) through complementary base pairing (Figure 7B).

When observing the ribosome pairs participating in this type of interaction directly in the tomogram after the subtomogram averages of the elongation states were mapped back into the tomograms, we found that the density representing the rRNA expansion segments in the ribosome pairs never connects fully, and a gap between the two opposed ES20La and ES31Lb extensions exists (Figure 7A). To gain further details on the interactions and to structurally determine whether the expansion segments were forming inter-ribosomal kissing-loop interactions, we attempted refinements that included both ribosomes in the pair. To this end, we selected ribosome pairs comprising two ribosomes in the rotated 2 state, since this was the most abundant pair type. In these reconstructions, one of the ribosomes refined to high resolution, whereas the second remained poorly resolved (Figure 7C, left panel), indicating that its orientation relative to the first ribosome is variable. Subsequent 3D classification allowed us to separate up to three different disome classes, in which the relative orientation of the second ribosome relative to the first one in the pair changes (Figure 7C, right panels). In the three classes, the two densities representing the ES31Lb expansions formed a direct connection, while the ES20La densities were disconnected (Figure 7C, right panels). Multibody refinement (41) also showed, through one of the main components, the rotational movement of the second ribosome relative to the first one in the pair (Supplementary Movie S1). Consistent with the 3D classification, the two densities representing the ES31Lb expansions formed a direct connection through the entire motion, and a ES31Lb: ES20La interaction only briefly occurs. This is consistent with the stronger and more stable kissing-loop interaction possible between the ES31Lb loops, which can involve up to six base pairs. By contrast, the ES20La loop sequence, GCGA, can support only a much weaker interaction of two base pairs with the corresponding ES20La loop in the partner ribosome (Figure 7B). The weaker, and likely only transient, interactions mediated by the ES20La expansion likely provide the flexibility for the two interacting ribosomes to adopt variable orientations relative to one another.

Interestingly, this was the only type of interaction that was observed repeatedly. Although we identified other ribosomes within the clusters that were close enough to ribosomes from different polysomes to potentially form contact interfaces, the structural motifs involved varied from case to case. This variability suggests that these contacts most likely reflect incidental proximity rather than specific interactions that contribute to cluster stability. Overall, our analysis indicates that neighboring ribosomes from different polysomes form recurrent interactions mediated by rRNA expansion segments, which likely help maintain the structural integrity of these dense ribosome clusters.

### Ribosomes in the same polysome within dense ribosomal clusters adopt non-canonical colliding interfaces

Previous studies in non-neuronal cells have found that when two ribosomes translating the same mRNA collide within a polysome, they form a disome, creating a well-characterized top-top 40S-40S interaction interface (23,59–62). In these collisions, the leading ribosome, located closer to the 3ʹ end of the mRNA, stalls first in an unrotated state containing a P-site tRNA, and then the trailing ribosome collides with the stalled ribosome and becomes trapped in the rotated 2 state, forming a disome. The interface between the two ribosomes is recognized by collision factors and serves as the primary substrate for pathways that resolve translational stalling.

Across all 341 tomograms, our analysis of ribosomes translating the same mRNA showed that many were spaced far enough apart from their upstream or downstream neighbors to avoid direct interaction. However, 12% of ribosomes were in close contact with adjacent ribosomes on the same mRNA (Figure 8A). In this analysis, and consistent with previous work (57), we classified ribosomes as translating the same mRNA and positioned to form a collision interface when the particle-to-particle distance was 150–270 Å and the distance between the mRNA exit site of one ribosome and the mRNA entry site of the neighboring ribosome was 85 Å. In some cases, only two ribosomes were in contact, forming a disome (e.g., cluster 7 in Supplementary Figure S10A). In other cases, longer arrays were observed, with 10 or more ribosomes contacting one another along the same mRNA (e.g., cluster 8 in Supplementary Figure S10A).

**Figure 8.**
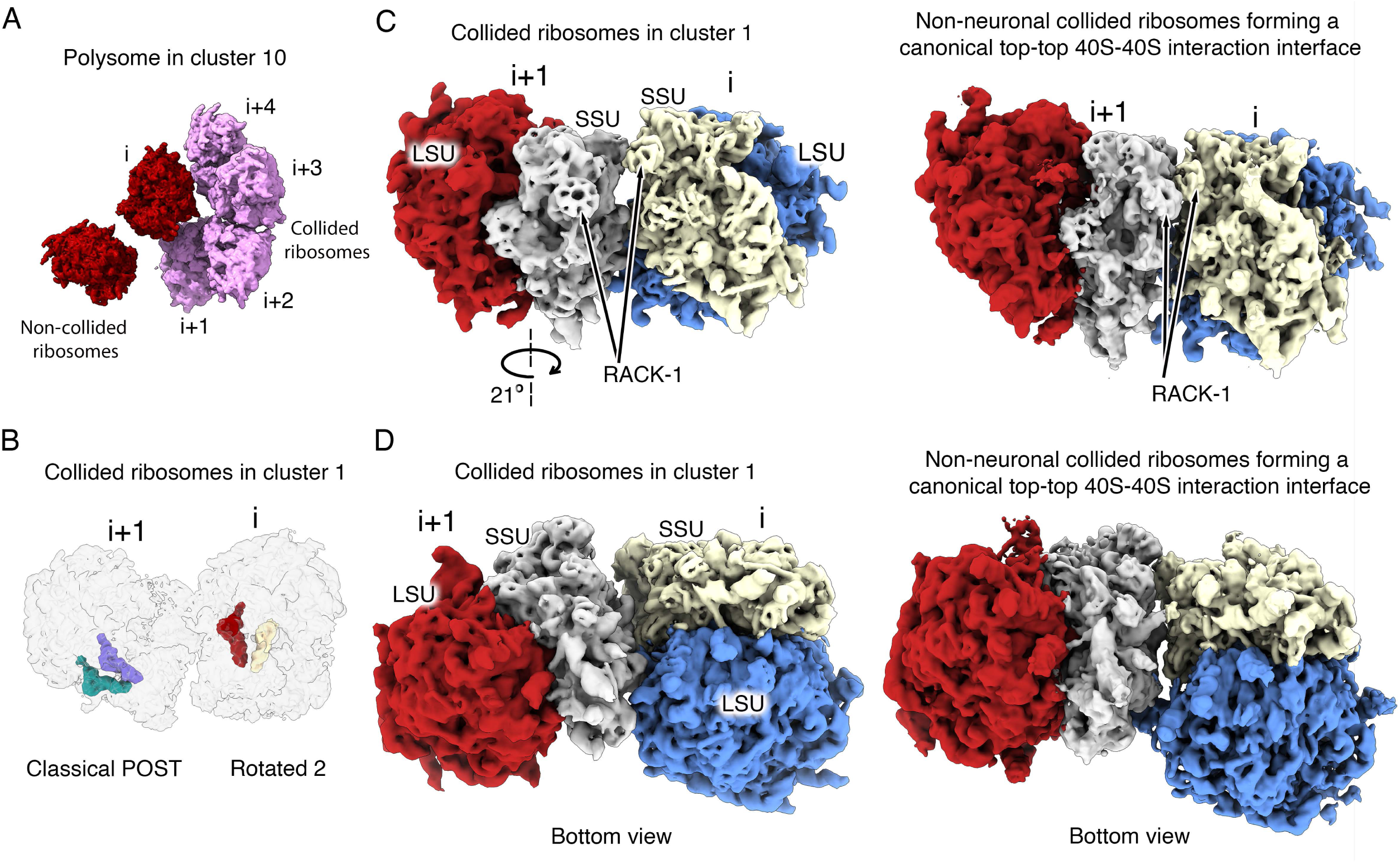
Interactions between ribosomes on the same mRNA molecule within ribosome clusters. (A) Elongation intermediates in cluster 10 illustrate that ribosomes within the same cluster can either remain isolated (non-collided) or form long arrays of ribosomes that interact with one another (collided), creating novel collision interfaces not previously described. The leading (i) and trailing (i+x) ribosomes in the collided stretch are labeled. (B) Semitransparent view of the cryo-EM map of the two collided ribosomes in cluster 1, showing their tRNA molecules, which define the elongation state of these ribosomes. The leading and trailing ribosomes in the disome are labeled as ‘i’ and ‘i+1’, respectively. (C) Side view of the surface representation of the cryo-EM map for the two collided ribosomes in cluster 1 (left) to visualize the interaction between the 40S subunits in the two collided ribosomes. An equivalent view is shown in the right panel for comparison with two ribosomes forming a canonical top-to-top 40S–40S collision in non-neuronal cells. The cryo-EM map for this disome was created using entries EMDB-0195 and EMDB-0197. The arrow diagram in the left panel indicates that the trailing ribosome in this disome is rotated by 21° relative to the trailing ribosome in the non-neuronal collision. The rotation axis is also indicated. Leading and trailing ribosomes are labeled as in (B). The large (LSU) and small (SSU) subunits of the ribosomes are labeled. The position of the RACK-1 protein is also indicated. (D) Bottom view of the same disome cryo-EM maps showing how the two ribosomes interact in the two types of collisions.

Most ribosomes within these disomes and longer arrays in the same mRNA were in the rotated 2 state (79%), while the remainder were in the rotated 1 (4%), classical PRE (8%), or classical POST (9%) states. This distribution closely resembled that of the overall ribosome population across all tomograms (Supplementary Figure S9), suggesting that disomes and longer ribosome arrays within the same mRNA incorporate the different elongation intermediates in proportions similar to those of the total population.

To determine whether elongation intermediates within disomes of collided ribosomes followed a pattern similar to that described for canonical collisions (leading ribosome containing a P/P tRNA and a trailing ribosome in the rotated 2 state) (23,59–62), we analyzed the states of adjacent leading and trailing ribosomes. In 64% of cases in which the leading ribosome was in the rotated 2 state, the trailing ribosome was also in the rotated 2 state. Rotated 1, classical PRE, and classical POST (Figure 8B) states were also observed in the trailing position behind rotated 2 ribosomes, but much less frequently. In the remaining 36% of cases, the leading ribosome was in the rotated 1, classical PRE, or classical POST state, and the trailing ribosome could adopt any of the other elongation intermediates without a clear preference. These results indicate that, although rotated 2–rotated 2 pairs were the most common, the collided ribosomes did not follow the canonical collision pattern or any other strict pattern in terms of the elongation state of the leading or trailing ribosomes forming the disome.

Ribosomes at the 3ʹ-most end position of long arrays of collided ribosomes were of particular interest because, based on their position, they are the most likely to represent the initially stalled ribosomes, whereas the closely trailing ribosomes behind them are more likely to be simply collided. To determine whether a specific elongation intermediate was enriched at this leading position, we recorded the elongation state of the 3ʹ-most ribosome in 83 long arrays of collided ribosomes identified within 240 polysomes from 100 representative dense ribosome clusters (Supplementary Figure S10A). In most cases, the leading ribosome was in the rotated 2 (73%) or rotated 1 states (16%). Classical PRE and classical POST states also occupied this position, but only in 5% and 6% of cases, respectively. These results indicate that the rotated states tend to occupy the 3ʹ-most end position of the long arrays of collided ribosomes more frequently than would be expected from the percentages of these intermediates in the total population (Supplementary Figure S9).

Importantly, the ribosomes within these dimers or longer collided arrays do not adopt an interaction interface resembling the canonical top-to-top 40S–40S collision interface previously described (23,59–62). Instead, as illustrated for a representative disome from cluster 1, the trailing ribosome is rotated by approximately 21° relative to its position in the canonical top-to-top 40S– 40S collision interface (Figure 8C, left and right panels). Although the magnitude of this rotation varies among collided ribosome pairs, it generally reconfigures the collision interface. Rather than being formed primarily by structural elements on the solvent-facing surfaces of the 40S subunits of both the leading and trailing ribosomes (Figure 8D, right panel), the interface shifts to one in which both the 40S and 60S subunits of the leading ribosome contact mainly the 40S subunit of the trailing ribosome (Figure 8D, left panel). This rearrangement also separates the RACK1 proteins bound to the heads of the two 40S subunits, preventing the RACK1–RACK1 contact characteristic of the canonical collision interface (Figure 8C). Among the 240 polysomes contained in the 100 clusters analyzed, we did not observe any disome interface with ribosomal orientation and distances that correspond to a canonical top-to-top 40S–40S collision.

In summary, the study of the ribosomes that stall and collide while reading the same mRNA in the dense ribosome clusters, revealed that they interact with each other by creating interfaces that are always different from the canonical top-to-top 40S–40S interfaces described between collided ribosomes in non-neuronal cells, therefore preventing these associations from being recognized by the collision factors that resolve canonical translational stalling and that dissociate the ribosomal subunits and initiate the degradation of the mRNA and nascent peptide.

## DISCUSSION

### A large proportion of ribosomes in P5 neurons are stalled

Earlier analyses established that the sedimented ribosome clusters analyzed in this study are a ribosomal fraction found in neurons and are distinct from the P-bodies and stress granules involved in RNA trafficking and storage (63). In particular, previous proteomic analysis of neuronal RNA granules revealed a lack of proteins associated with translating polysomes (eEF2) or P-body proteins (decapping enzymes) (14,47,64). Instead, some of the RBPs found in stress granules were also identified in neuronal RNA granules (FMRP, G3BP1). However, the absence of free 40S subunits (a typical component of stress granules) in the neuronal RNA granules (20) indicated that stress granules are not present in the pellet fraction containing the sedimented ribosome clusters.

Our negative-staining electron microscopy analysis of the ribosome clusters in the pellet fraction and those contained in fractions 5/6 and 2/3 (equivalent to polysomes and monosomes in ribosome purification in other cell types) concluded that the overall morphology of the ribosome clusters included in all these fractions was similar (Supplementary Figure S1A & S1B). More importantly, they were markedly different from the linear array of ribosomes forming the polysomes purified from the liver of the same animals (Supplementary Figure S1D). Subsequent single-particle cryo-EM analysis showed that about half of the elongation intermediates included, both in the ribosome clusters in the pellet fraction and in fractions 5/6 and 2/3, were stalled. Given the different sedimentation behavior of the ribosome clusters migrating to the pellet and those migrating in fractions 5/6 and 2/3, our study suggests the possibility that the ribosome clusters in fractions 5/6 and 2/3 serve as precursors for the formation of the dense ribosome clusters in the pellet fraction through an still undescribed compaction mechanism that eventually converts them into denser clusters.

### Sequence elements underlying stalling

Cryo-EM and sequence analysis of stalling sites independently predict that the mRNA adopts a secondary structure close to its entry site on the ribosome. Both the enrichment for Gs (Figure 5B, middle panel) and the density observed in the cryo-EM maps at the entrance of the mRNA channel (Figure 5C) are consistent with these sequences forming a G-quadruplex structure. However, the local resolution of our maps in this region is not sufficient to unambiguously model this part of the structure, thus other alternative secondary structures cannot be ruled out. Another independent line of evidence consistent with secondary structure at the mRNA entrance site is the size of RPFs generated from these ribosomes. Our initial study found RPFs considerably longer (33-37 bp) than normal (28-30 bp) from pellet and fractions 5/6 derived ribosomes, and the increased length was at the 3’ end of the RPFs near the RNA entrance site (20). More recently, we compared different nuclease concentrations and replicated this result using the nuclease concentration from the previous study; however, higher concentrations of nuclease generated RPFs of normal size (55). These results are consistent with the mRNA adopting a secondary structure near its entry site into the ribosome, thereby requiring higher nuclease concentrations for cleavage.

In conjunction with the mRNA secondary structure, we found enrichment of several amino acids in nascent peptides from these ribosomes (Figure 5B, middle panel). A previous observation also suggests that the nascent peptide within the exit tunnel may still contribute to stalling. In a previous cryo-EM study, ribosomes from the pellet fraction were treated with puromycin (48). In non-neuronal ribosomes, puromycin treatment typically results in the release of the puromycylated nascent peptide from the ribosome (65,66). In contrast, in neuronal ribosomes from the pellet fraction, the puromycylated nascent peptide remained within the exit tunnel of the large subunit (48), implying the existence of interactions between the nascent peptide and the exit tunnel. However, the amino acids that are enriched in the nascent peptides in the stall ribosomes span a broad range of physicochemical classes, including acidic, basic, hydrophobic, polar, and neutral. Therefore, it is difficult to imagine, from the existing data, a mechanism by which they might contribute to stalling. Further studies are needed to clarify this aspect of the stalling mechanism.

Taken together, these findings indicate that the stalling mechanism is likely multifactorial. In addition to possible contributions from mRNA sequence or structure, and amino acids at the beginning of the peptide tunnel, other factors are also likely to be involved. One such factor may be the lower levels of eEF2 in neurons (67), which could favor accumulation of ribosomes in rotated states, as eEF2 is required for the 40S subunit to backrotate from the rotated states into the classical POST state.

Finally, FMRP, an RNA-binding protein in the pellet fraction (Figure 4) (20), is also known to bind G-quadruplex structures and kissing-loop interactions (68). This raised the possibility that FMRP might contribute directly to ribosome stalling. However, loss of FMRP does not alter either the positions at which ribosomes stall or the relative abundance of stalled ribosomes (55), arguing against a direct role in the stalling mechanism itself. Instead, if G-quadruplexes and kissing-loop interactions are present within these ribosomal clusters, they may help explain why FMRP is enriched at these sites. Based on these findings, our current model is that the initial stalling event is triggered by features of the mRNA sequence and possibly amino acids in the beginning of the peptide tunnel. However, the longer-term stabilization of stalled ribosomes into clusters is driven by interactions among ribosomes within the clusters, as observed by cryo-ET in this study (Figures 7 and 8). The compaction of these clusters likely also depends on additional nervous system-specific factors and interactions that remain to be identified.

### Importance of polyamines in preserving elongation intermediates in the neuronal RNA granules

Two previous studies conducted cryo-EM analysis and image classification of ribosomes incorporated into neuronal RNA granules (20,47). Both studies found that most ribosomes were in the rotated 2 state. Interestingly, neither study reported the high proportions of classical PRE or POST elongation intermediates observed in this study across samples from pellet, polysomes, and monosomes. One factor that may be causing these differences is recent improvements in cryo-EM image classification tools (69), which have enhanced the ability to distinguish elongation intermediates.

However, the improved buffer conditions used in this study likely better preserved the elongation states present in the dense ribosome clusters from the pellet fraction. In particular, the buffers in our study incorporate polyamines (0.04 mM spermine and 0.5 mM spermidine). Cryo-EM structures of the ribosome obtained from cellular lamellas cut with a cryo-focused ion beam revealed that each ribosome, in its unperturbed cellular environment, can bind up to 14 polyamine molecules, contributing to the structural stability of the regions near their binding sites (70).

There is also abundant evidence indicating that polyamines stabilize tRNA-ribosome interactions (71), accelerate codon recognition, and modulate translation (72). This protective role of polyamines was likely essential for observing additional elongation intermediates in our study, particularly those loaded with an E-site tRNA, which binds with the lowest affinity (73,74). Finally, consistent with a stabilizing role of spermidine in the structural stability of the small subunit (75), we found that including polyamines in the buffers for this study enabled us to achieve higher local resolution for the small subunits in the cryo-EM maps presented here compared to previous structures of ribosomes included in neuronal RNA granules (20,47).

### Ribosomal RNA expansion segments mediate interactions between ribosomes in the neuronal RNA granules

rRNA expansion segments are peripheral structural elements located on the solvent-exposed surfaces of both the large and small ribosomal subunits. Given their position, they are unlikely to contribute directly to the ribosome’s core catalytic functions, such as peptidyl transferase activity and decoding. Instead, current evidence suggests that these extensions primarily serve as interaction platforms for eukaryote-specific ribosomal proteins, translation regulators, and factors involved in ribosome biogenesis and quality control (76–78). Nevertheless, their precise roles remain poorly understood.

A recent study (58) reported that, under nutrient-deprivation stress, ribosomes in rat hippocampal neurons form disomes through interactions mediated by rRNA expansion segments. In these disomes, the two ribosomes contact each other via their large subunits, specifically through a homotypic interaction between the ES31Lb segments of the neighboring ribosomes. The two ES31Lb segments form a “kissing-loop” interaction formed by unpaired complementary nucleotides at the tips of the two ES31Lb elements. Similarly, another study (79) found that puromycin treatment induces the formation of reversible disomes in primary cultures of rat hippocampal neurons. In that case, disome stabilization was mediated either by the same homotypic ES31Lb– ES31Lb RNA–RNA interaction or by a heterotypic interaction between ES20La in one ribosome and ES30La in the other.

Although the nutrient-deprivation-induced (58) and puromycin-induced (79) disomes involving ES31Lb and ES20La resemble the interactions observed in the ribosome clusters from the pellet fraction in our study, they differ in several fundamental ways. In the two described cases, these interactions stabilized pairs of otherwise isolated ribosomes, resulting in disome formation. In addition, the ribosomes forming disomes in nutrient-deprived neurons (58) were largely empty and lacked mRNA and tRNAs, whereas those in puromycin-treated neurons (79) were either idle or assembled into non-translating complexes associated with eIF5A, SERBP1, and eEF2. By contrast, in the ribosome clusters from the pellet fraction in our study, ES31Lb and ES20La link ribosomes within long polysomes containing multiple ribosomes stalled during elongation.

The ES31Lb– and ES20La-mediated interactions observed in the ribosome clusters from the pellet fraction in our study also differ in several important aspects from those described for disomes of inactive or hibernating ribosomes in neurons (58). Experiments showed that the ES31Lb-mediated interaction is both necessary and sufficient for disome stability. This interaction also requires magnesium concentrations above 12.5 mM to remain stable. Indeed, the model for the formation of disomes under stress was linked to the increased intracellular magnesium concentrations achieved during stress (58). However, previous structural studies from our own group found that the ribosome clusters in the pellet fraction were stable at a magnesium concentration of 2.5 mM (20). Comparisons of ES21Lb sequences across species suggest it can form in 17% of chordate species, including humans. However, mouse (*Mus musculus*) ribosomes do not dimerize through this mechanism. Surprisingly, a study analyzing the role of the FMRP protein in regulating ribosome clusters found that these clusters can also be purified from mouse brains using the same approach we used in our study. The ribosome clusters in the mouse pellet fraction had a similar overall morphology in negative-staining electron micrographs to those found in the rat (55). However, this study did not provide a cryo-ET analysis to determine whether any interactions among ribosomes within the mouse clusters were mediated by ES31Lb or other rRNA extensions. These differences suggest that the interactions mediated by the ES31Lb and ES20La segments in the dense ribosome clusters observed in the pellet fraction of our study are probably not the only ones mediating their clustering, and that other interactions are likely required for the full stability of these structures.

It is also possible that multiple RBPs contribute to the stability of the dense ribosomal clusters in the pellet fraction in our study. However, no obvious protein densities were visible in our tomograms. It is possible that proteins involved in these ribosome interactions stabilize kissing-loop interactions or facilitate liquid-liquid phase separation, contributing to the assembly of the ribosome clusters. Most phase-separating proteins contain large intrinsically disordered regions, or fluctuate among many conformations (80,81). This conformational heterogeneity makes them difficult to resolve by cryo-ET. Several of the proteins identified as part of the dense ribosome clusters from the pellet fraction through previous proteomic analysis (14,47,64) and in this work (Figure 4), including FMRP, Staufen, and G3BP1 have also been found in neuronal biomolecular condensates, such as transport granules, stress granules, and synaptic condensates. Nevertheless, a limitation of our study is that it does not elucidate how these proteins may contribute to stabilizing the observed ribosomal interactions within these clusters. Additional studies will be necessary to clarify the role of phase-separation proteins in the mechanism of assembly of the dense ribosome clusters in the pellet fraction.

### Collided ribosomes included in the neuronal RNA granules do not trigger the ribosome-associated quality control response

Translation elongation occasionally stalls in cells due to oxidized or truncated mRNA or higher-order mRNA structures (82,83). In addition, proline-rich and lysine-rich sequences can cause ribosome stalling, but for different mechanistic reasons. Proline is a poor peptidyl-transfer substrate, and its rigid ring makes peptide bond formation difficult (84,85). Poly-lysine can also stall ribosomes because the positively charged lysine chains interact with the exit peptide tunnel, which is negatively charged, and slow down elongation (86). In most cells, once a ribosome has stalled at a specific sequence, the ribosome following it in the polysome will only elongate until it collides with the stalled one. This collision activates the RQC response, resulting in the recycling of the 40S and 60S subunits and degradation of the arrested nascent protein. It also initiates a process called NO-GO decay, which leads to destruction of the mRNA (87,88). A critical step in activating these responses is the assembly of a collided disome structure. This disome is made of the stalled ribosome and the next ribosome in the polysome colliding into it. The paused stalled leading ribosome contains a P/P tRNA, and the trailing collided ribosome contains A/P and P/E tRNAs (23). This complex is not simply two juxtaposed ribosomes. The 40S-40S interaction interface formed by the two ribosomes serves as a defined platform recognized by various collision-specific factors that trigger the RQC and NO-GO decay responses (82,83).

Our cryo-ET analysis revealed that triggering of the RQC and NO-GO decay responses is avoided in neurons through the unique three-dimensional organization of these dense ribosome clusters in the pellet fraction. First, we found that both elongation intermediates included in the disomes in the ribosome clusters in the pellet fraction are mainly in a rotated 2 state (Supplementary Figure S10). This is drastically different from the fixed pattern of canonical collided disomes that typically trigger the RQC and NO-GO decay responses, and in which the leading and trailing collided ribosomes are in a classical POST and rotate 2 states, respectively. Furthermore, the interactions observed between the ribosomes included in the dense ribosome clusters from the pellet fraction are also different from the 40S-40S interacting interface created by the two ribosomes in the canonical disome capable of triggering the RQC and NO-GO decay responses. Consequently, the ribosome clusters formed in neurons go undetected and do not trigger the RQC and NO-GO decay responses.

## DATA AVAILABILITY

The cryo-EM maps obtained in this study and the derived molecular models have been deposited in the Electron Microscopy Data Bank (EMDB) and the Protein Data Bank (PDB) with accession codes detailed in Supplementary Tables S1 to S7.

## SUPPLEMENTARY MATERIAL

Supplementary Data are available online.

## ACKNOWLEDGEMENTS

We thank staff members of the Facility for Electron Microscopy Research (FEMR) at McGill University for help with microscope operation and data collection.

## SUPPLEMENTARY FIGURES

**Supplementary Figure S1.**
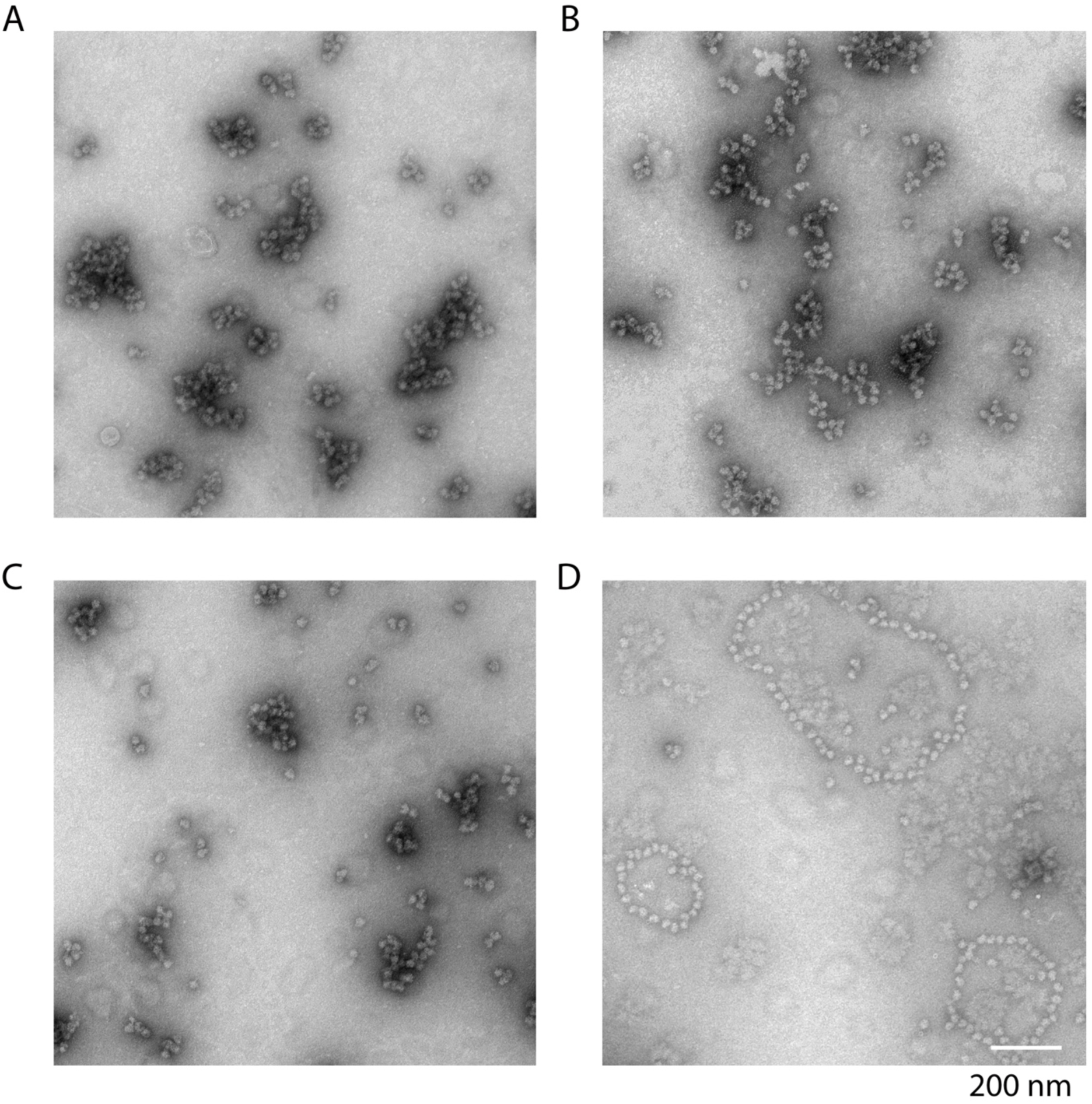
Negative staining electron micrographs of neuronal and liver ribosomes. (A) Negative staining electron microscopy images of neuronal RNA granules (dense ribosome clusters from the pellet fraction) purified from P5 rat brains (pellet fractions). These are the same samples that were imaged by cryo-electron tomography in this study. (B) Negative staining electron microscopy images of samples from fractions 5-6 purified from P5 rat brains. (C) The electron microscopy images show samples from fractions 2-3 purified from rat brains under negative staining. (D) Negative staining electron microscopy images of polysomes (fractions 5-6) purified from P5 rat livers.

**Supplementary Figure S2.**
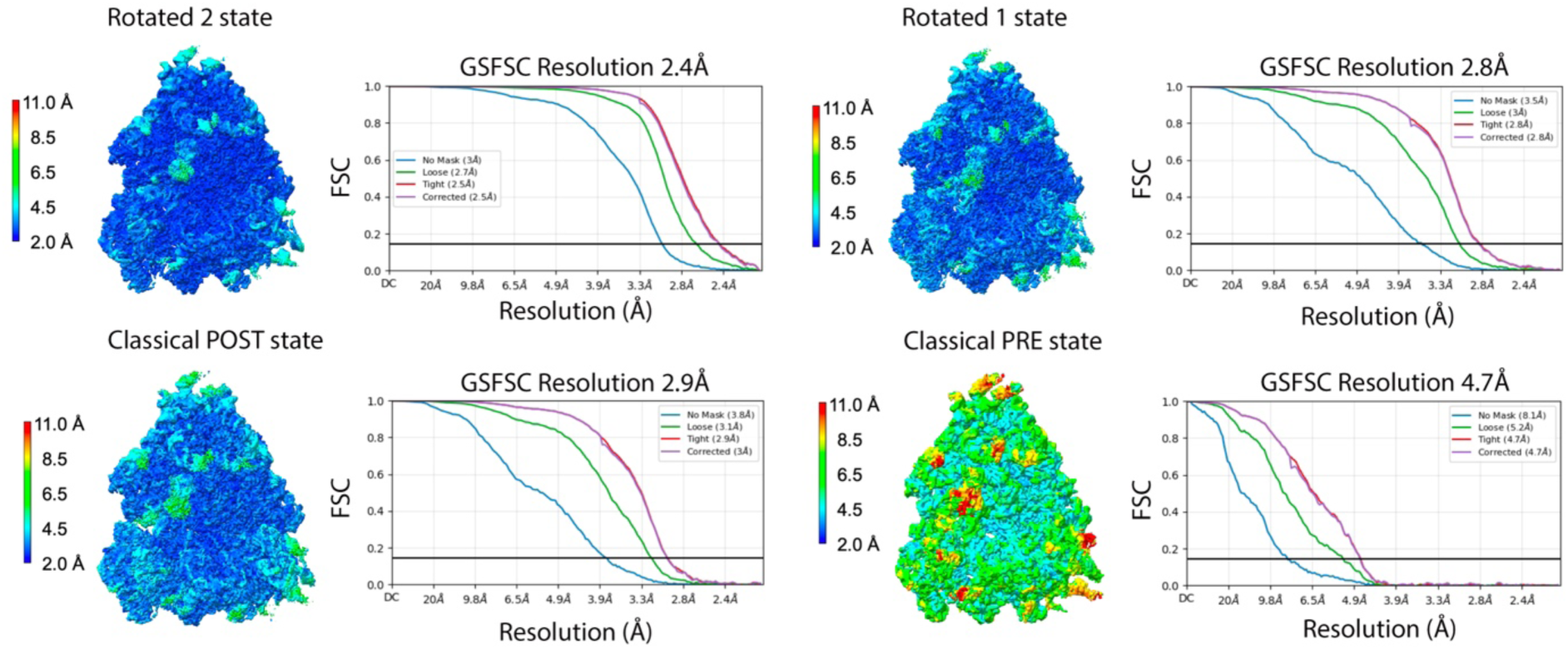
Local resolution analysis of the cryo-EM maps of the elongation intermediates in the purified dense ribosome clusters from the pellet fraction. The cryo-EM density maps of the elongation intermediates within the dense ribosome clusters from the pellet fraction are colored by local resolution. The elongation intermediate identity is indicated on top of each map. The Gold-Standard Fourier shell correlation (GSFSC) plots for each map are shown. The FSC curves were calculated using different masks. In the ‘No Mask’ plot, the raw FSC curve was calculated between two independent half-maps without masking. Loose’: FSC calculated after applying a loose soft solvent mask to both half maps. The loose mask is calculated by thresholding the density map at 50% of the maximum density value. The resulting volume is dilated to create a soft mask. Voxels in the mask within 25 angstroms of the threshold region receive a mask value of 1.0. Voxels between 25 and 40 angstroms fall off with a soft cosine edge, and voxels outside 40 angstroms receive a value of 0.0. ‘Tight’ is similar to the ‘Loose’ mask, except that the dilation distances are 6 angstroms for the value 1.0 distance and 12 angstroms for the value 0.0 distance. ‘Corrected’ is using the ‘Tight’ mask corrected by noise substitution. In this case, the two half maps have their phases randomized beyond a certain resolution, then the tight mask is applied to both, and a FSC is calculated. This FSC is used along with the original FSC before phase randomization to compute the corrected FSC. This accounts for correlation effects induced by masking. The resolution at which phase randomization begins is the resolution at which the no-mask FSC drops below the FSC = 0.143 criterion. The reported resolution is based on an FSC threshold of 0.143.

**Supplementary Figure S3.**
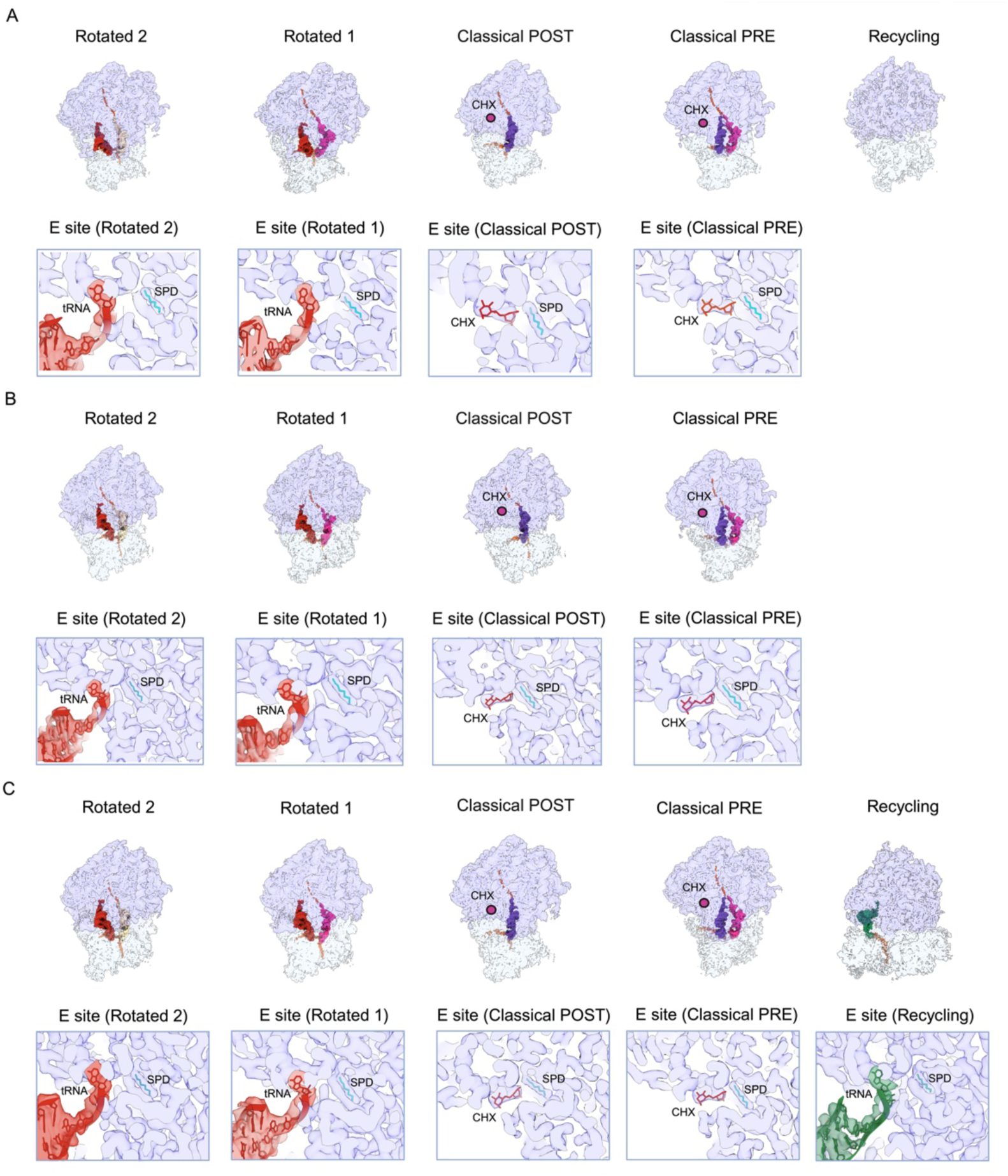
Elongation intermediates in cycloheximide-treated purified dense ribosome clusters from the pellet fraction and from fractions 5/6 and 2/3. (A) The top row shows the cryo-EM maps of elongation intermediates obtained through image classification from purified dense ribosome clusters contained in the pellet fraction after treatment with cyclohexamide. Densities representing the 40S and 60S subunits are displayed as transparent to highlight the tRNA molecules within the ribosome for each elongation state. Each elongation intermediate is characterized by the location of the tRNA in the A, P, and E sites. The bottom row shows the zoomed-in view of the density at the E site in cryo-EM maps of the classical PRE and classical POST elongation intermediates. The molecular models for cycloheximide (CHX) and spermidine (SPD) are shown docked into the density. It also shows the zoomed-in view of the density in the E site in the cryo-EM maps of the rotated 1 and rotated 2 elongation intermediates. The molecular models for the conserved CAA nucleotides at the 3ʹ end of E site tRNA molecule (in red) and for spermidine (SPD) are shown docked into the density. An equivalent panel to (A) showing the cryo-EM maps of elongation intermediates in the fractions 5/6 and 2/3 purified in the presence of cycloheximide is shown in (B) and (C), respectively. These panels also show the zoomed-in view of the density at the E site in these cryo-EM maps.

**Supplementary Figure S4.**
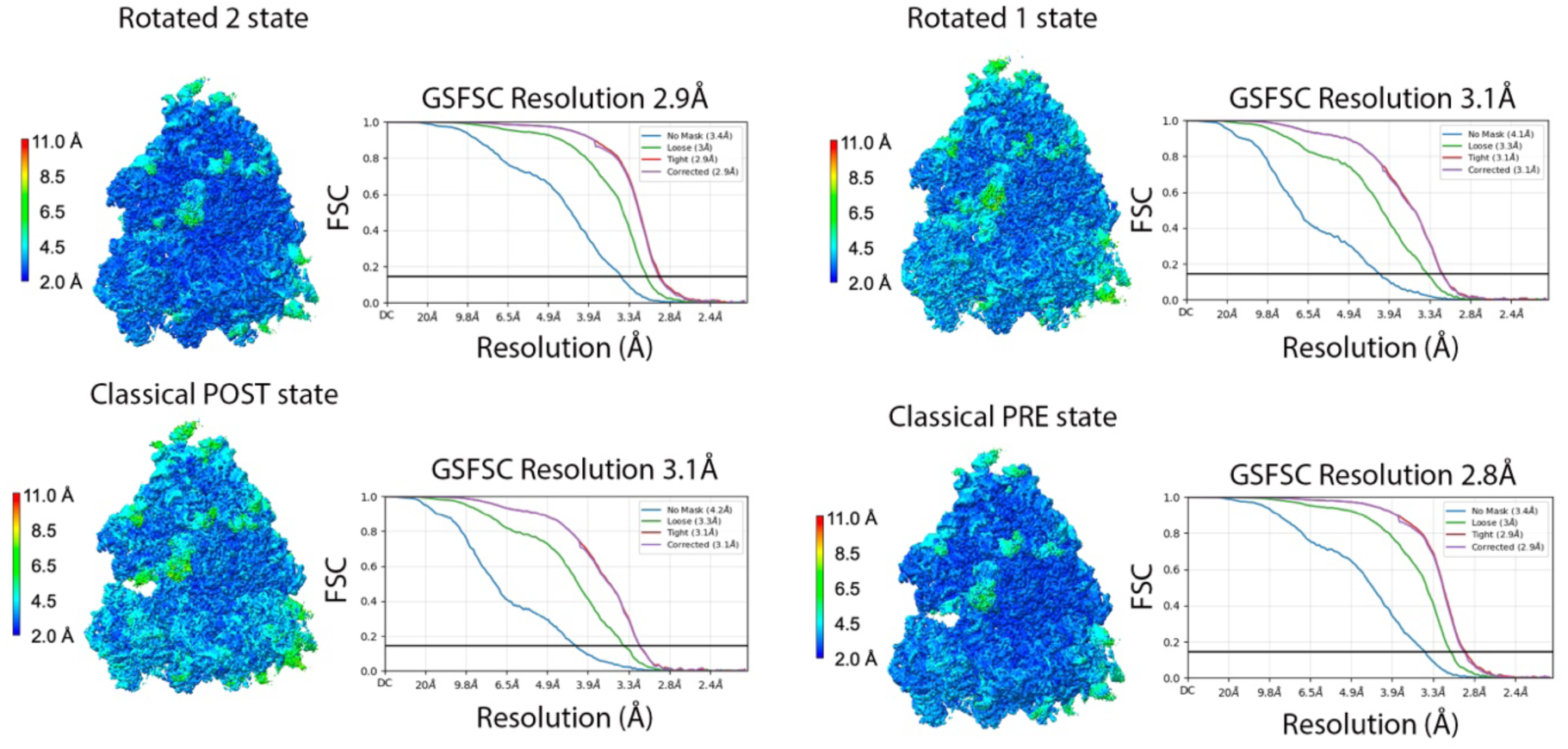
Local resolution analysis of the cryo-EM maps of the elongation intermediates in the cycloheximide-treated purified dense ribosome clusters from the pellet fraction. Cryo-EM density maps obtained for the elongation intermediates contained in cycloheximide-treated purified dense ribosome clusters from the pellet fraction are colored according to their local resolution. The elongation intermediate identity is indicated on top of each map. The graphs show the ‘No Mask,’ ‘Loose, ‘’ Tight, ‘and ‘Corrected’ FSC plots calculated as described in Supplementary Figure S2. The reported resolution is based on an FSC threshold of 0.143.

**Supplementary Figure S5.**
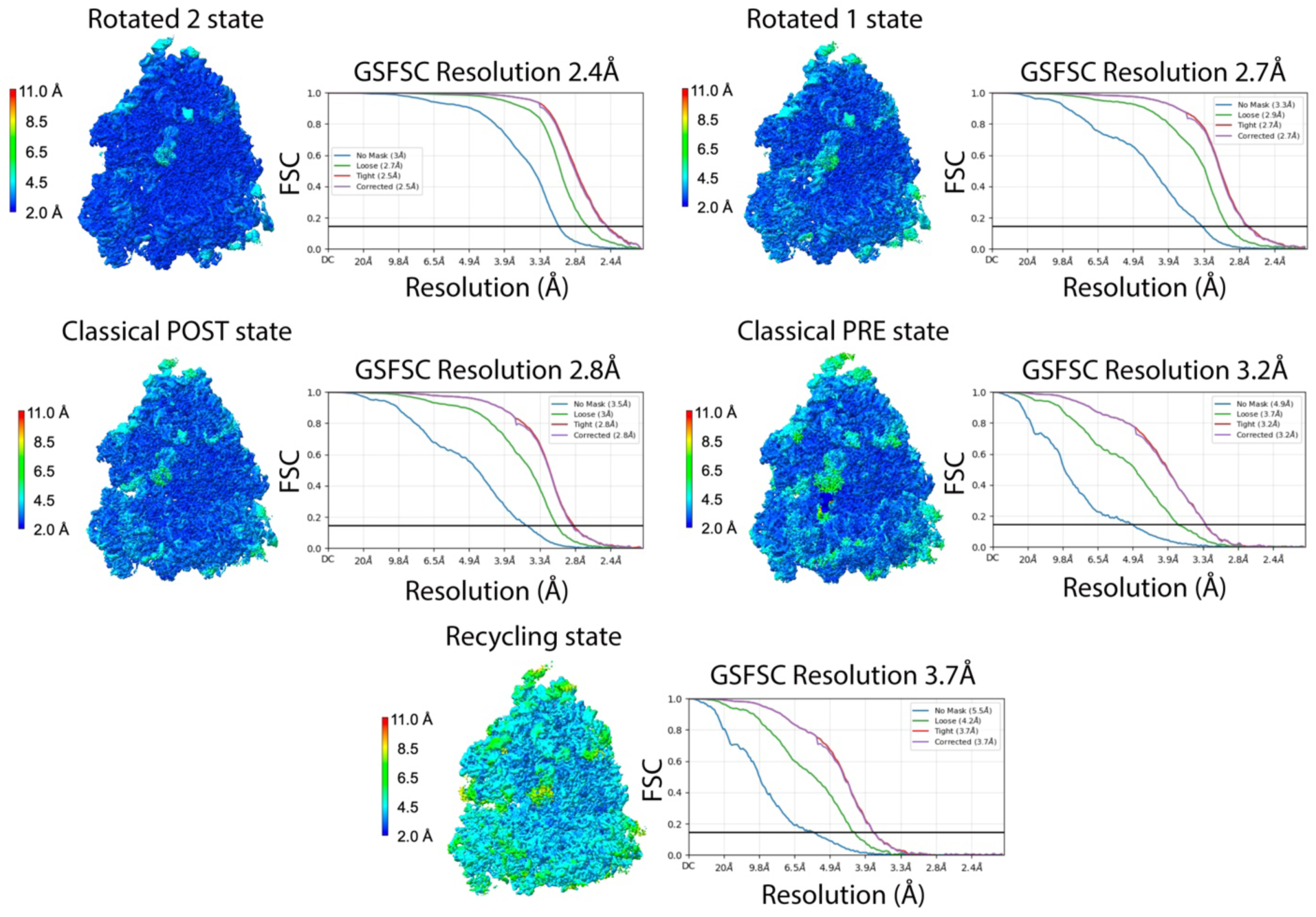
Local resolution analysis of the cryo-EM maps of the elongation intermediates from fractions 5/6. Cryo-EM density maps of the elongation intermediates in fractions 5/6 are colored by local resolution. The elongation intermediate identity is indicated on top of each map. The graphs show the ‘No Mask,’ ‘Loose, ‘’ Tight, ‘and ‘Corrected’ FSC plots calculated as described in Supplementary Figure S2. The reported resolution is based on an FSC threshold of 0.143.

**Supplementary Figure S6.**
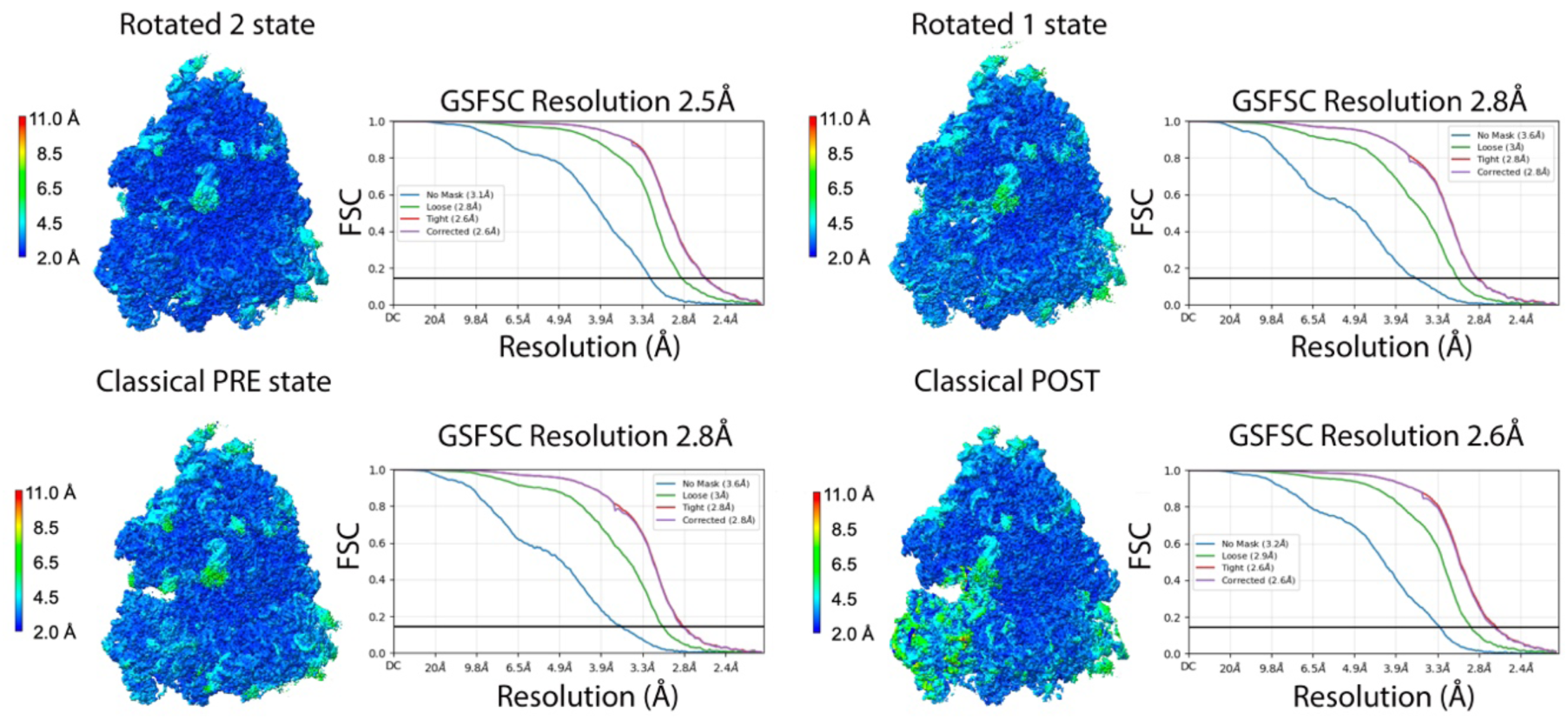
Local resolution analysis of the cryo-EM maps of the elongation intermediates from the cycloheximide-treated fractions 5/6. Cryo-EM density maps of the elongation intermediates contained in the cycloheximide-treated purified fractions 5/6 are colored by local resolution. The elongation intermediate identity is indicated on top of each map. The graphs show the ‘No Mask,’ ‘Loose, ‘’ Tight, ‘and ‘Corrected’ FSC plots calculated as described in Supplementary Figure S2. The reported resolution is based on an FSC threshold of 0.143.

**Supplementary Figure S7.**
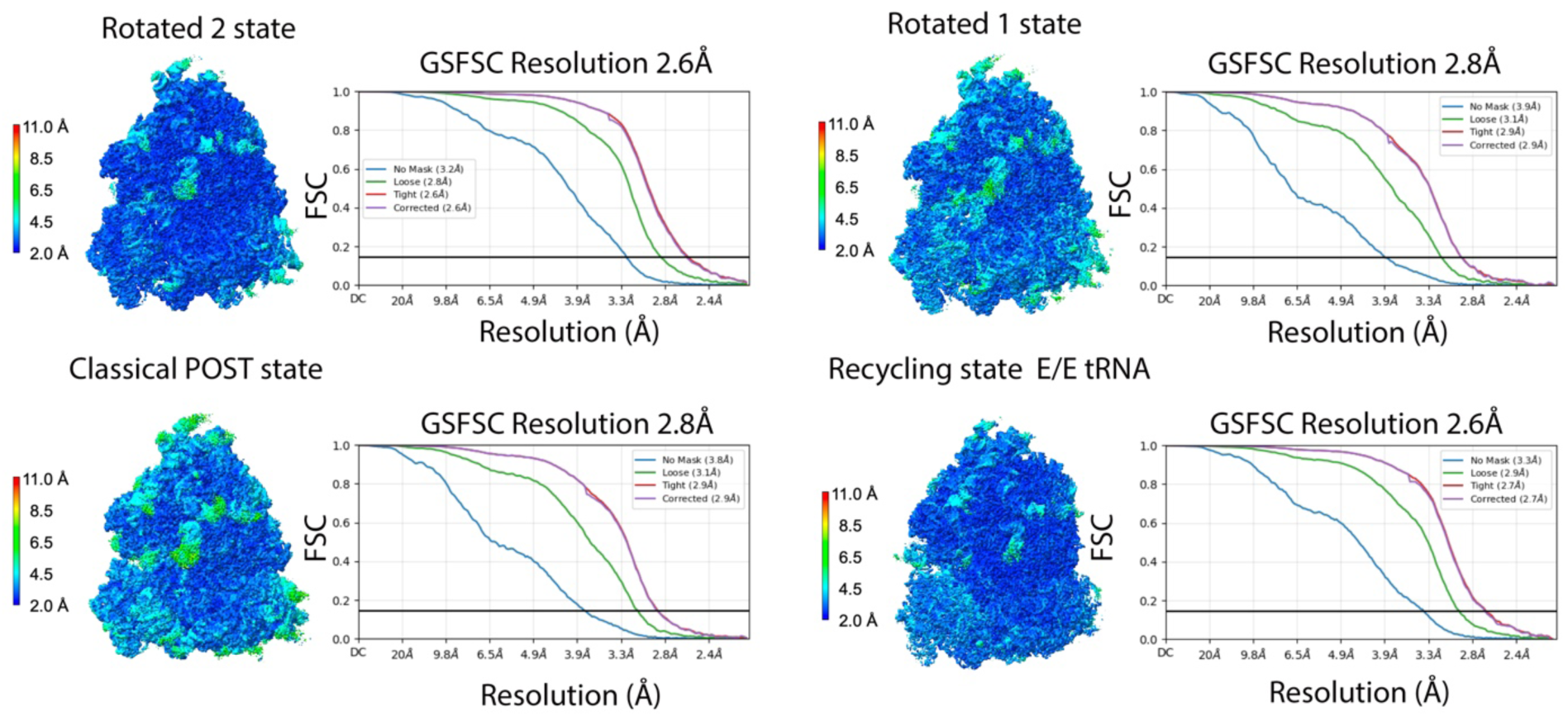
Local resolution analysis of the cryo-EM maps of the elongation intermediates in fractions 2/3. Cryo-EM density maps of the elongation intermediates in fractions 2/3 are colored by local resolution. The elongation intermediate identity is indicated on top of each map. The graphs show the ‘No Mask,’ ‘Loose, ‘’ Tight, ‘and ‘Corrected’ FSC plots calculated as described in Supplementary Figure S2. The reported resolution is based on an FSC threshold of 0.143.

**Supplementary Figure S8.**
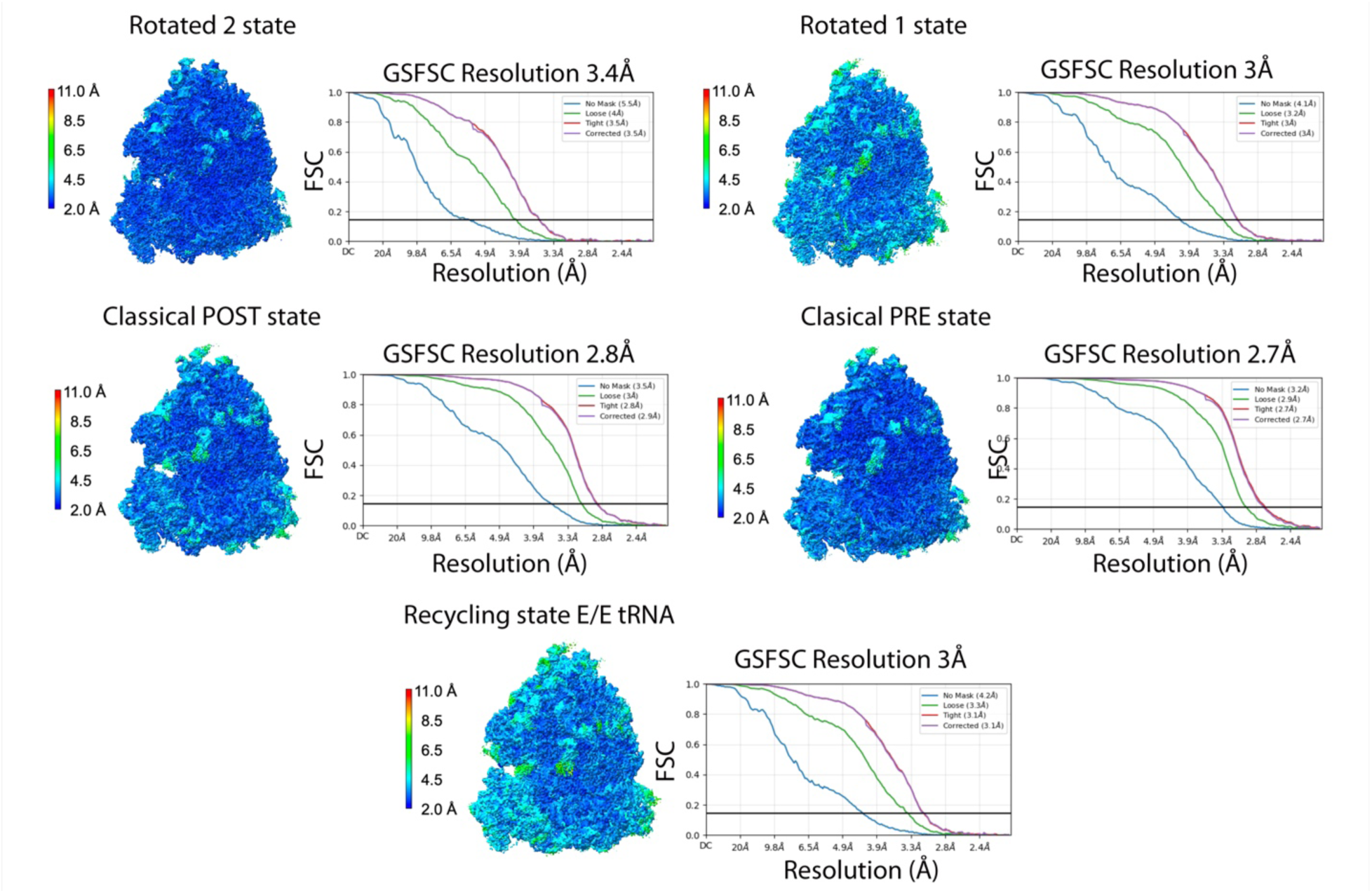
Local resolution analysis of the cryo-EM maps of the elongation intermediates in the cycloheximide-treated fractions 2/3. Cryo-EM density maps of the elongation intermediates in the cycloheximide-treated fractions 2/3 are colored by local resolution. The elongation intermediate identity is indicated on top of each map. The graphs show the ‘No Mask,’ ‘Loose, ‘’ Tight, ‘and ‘Corrected’ FSC plots calculated as described in Supplementary Figure S2. The reported resolution is based on an FSC threshold of 0.143.

**Supplementary Figure S9.**
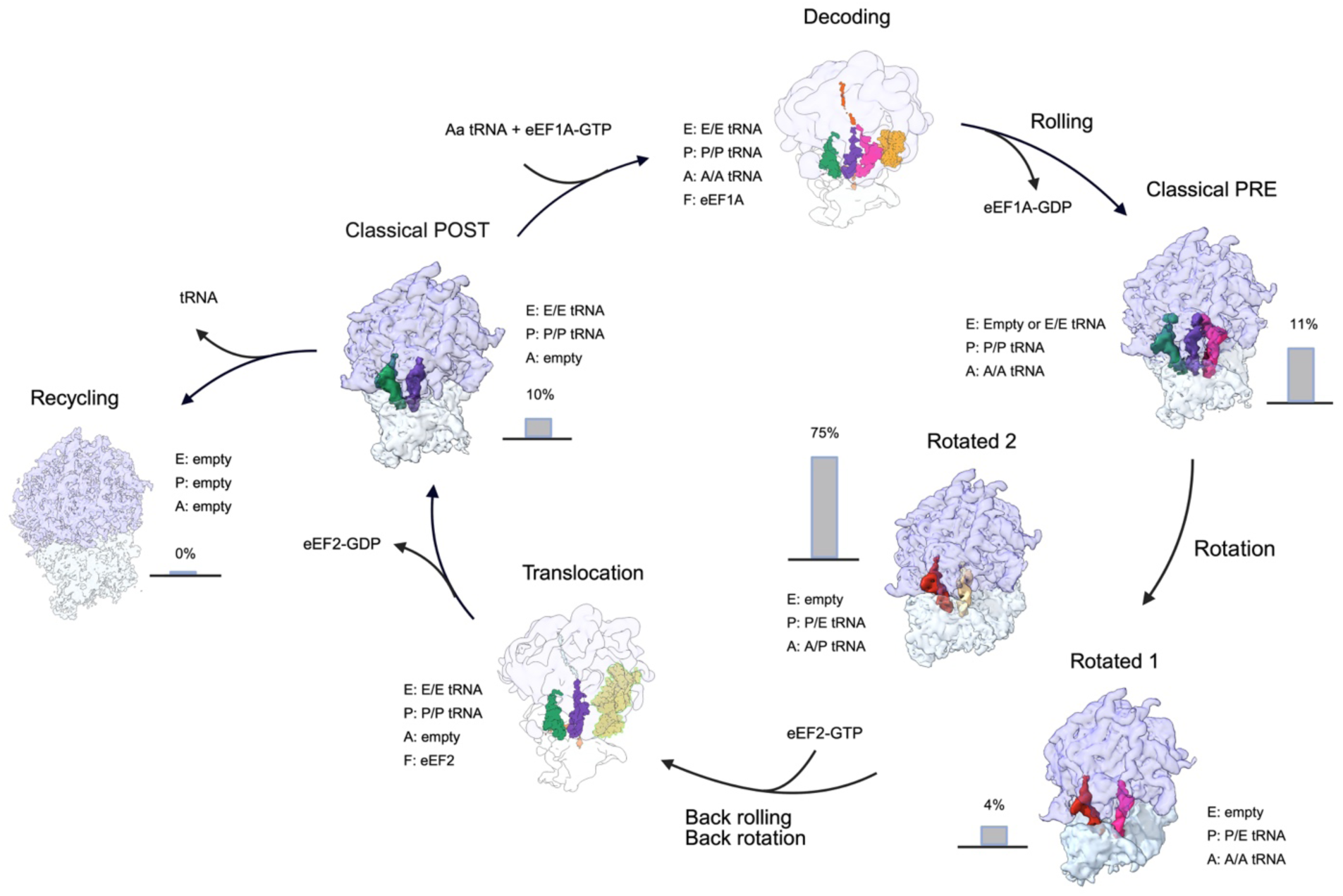
Elongation intermediates in purified dense ribosome clusters from the pellet fraction imaged by cryo-electron tomography. The subtomogram averages obtained from the tomograms of purified dense ribosome clusters from the pellet fraction are positioned within the canonical elongation cycle. The layout of this panel and labels are as in Figure 1A. The bar graph beside each map indicates the proportion of particles in that elongation state.

**Supplementary Figure S10.**
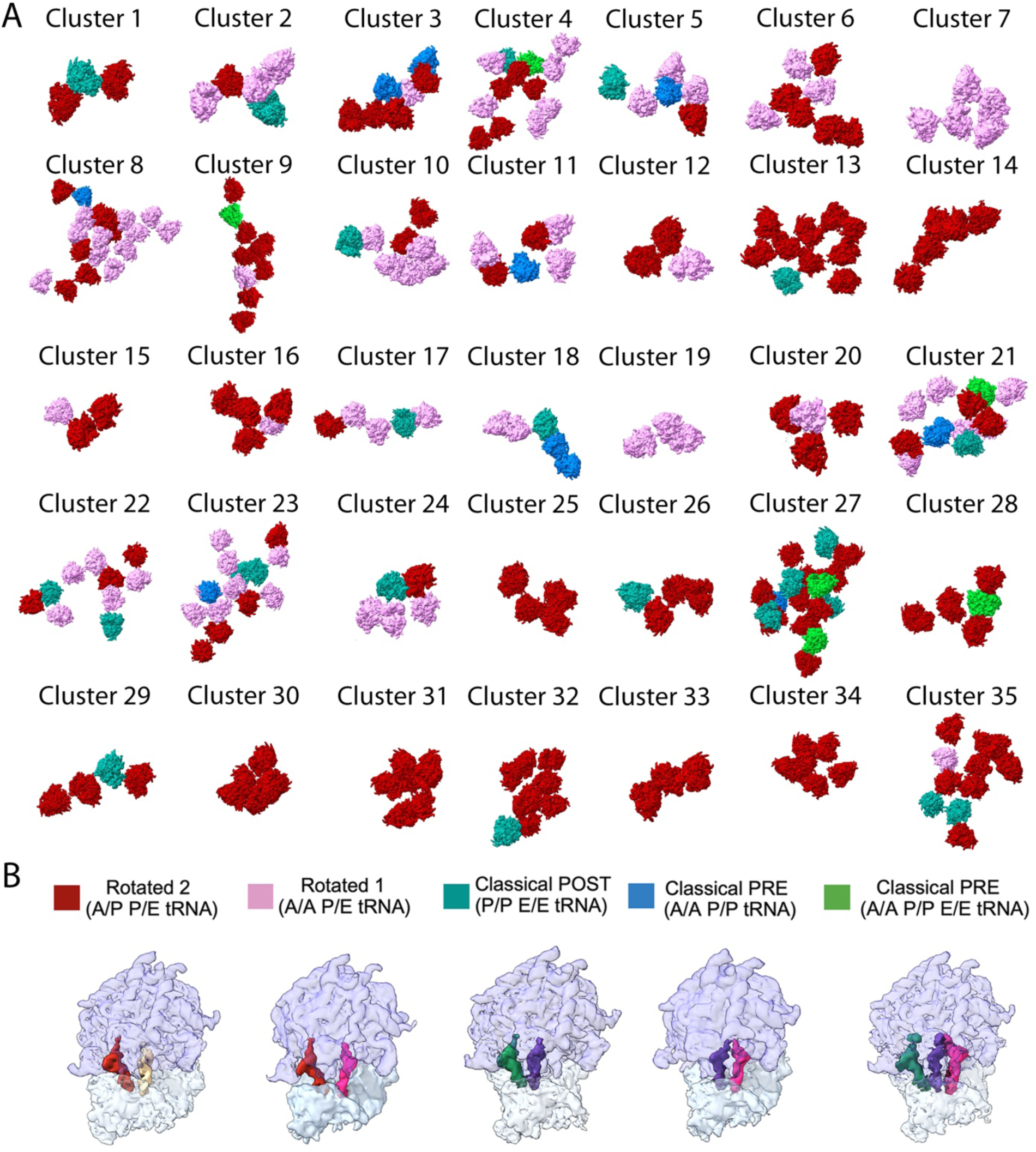
Ribosome clusters in purified dense ribosome clusters from the pellet fraction imaged by cryo-electron tomography. (A) Overview of 35 representative ribosome clusters out of the 100 randomly selected and analyzed from 30 cryo-tomograms obtained from purified neuronal RNA granules. These clusters were used to study the number of mRNA molecules and their trajectories within the clusters, as well as the multiple ways in which ribosomes interact with one another within the three-dimensional architecture of neuronal RNA granules. The color used to display each ribosome in the cluster identifies its elongation state, according to the color code indicated in panel (B).

## SUPPLEMENTARY TABLES

**Supplementary Table S1.**
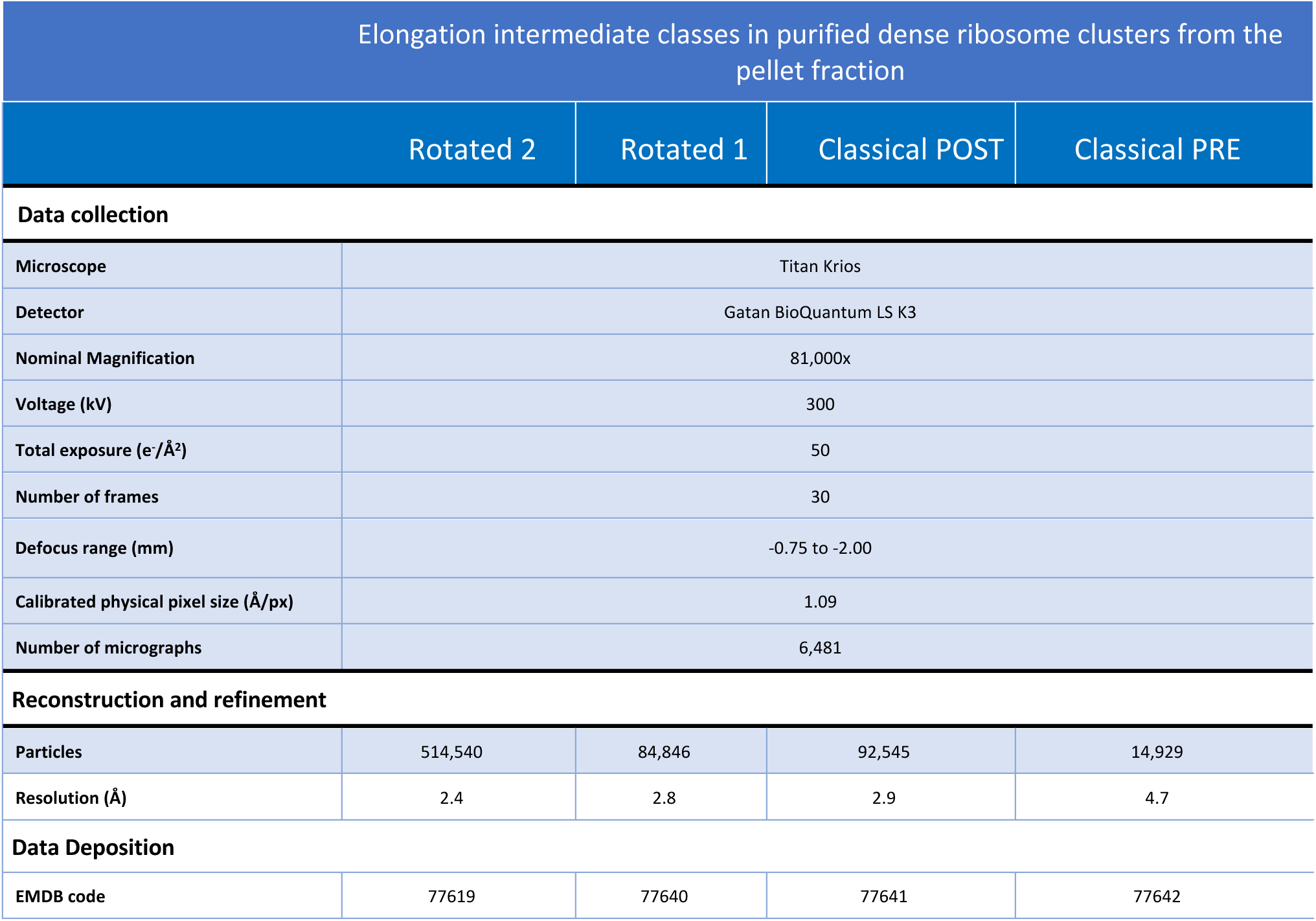
Cryo-EM analysis of the elongation intermediates in the purified dense ribosome clusters from the pellet fraction: data acquisition parameters and map statistics.

**Supplementary Table S2.**
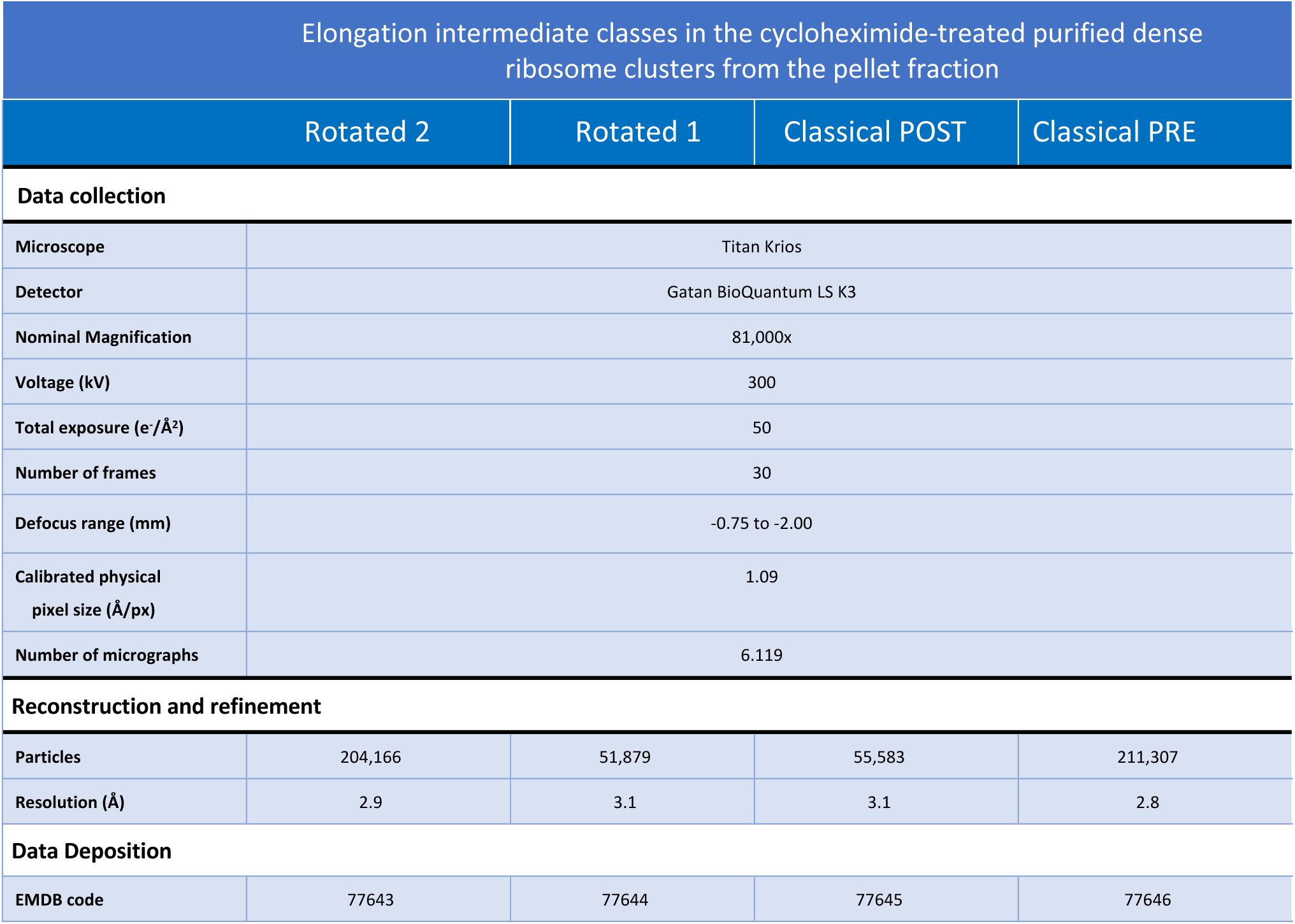
Cryo-EM analysis of the elongation intermediates in the purified cycloheximide-treated neuronal RNA granules: data acquisition parameters and map statistics.

**Supplementary Table S3.**
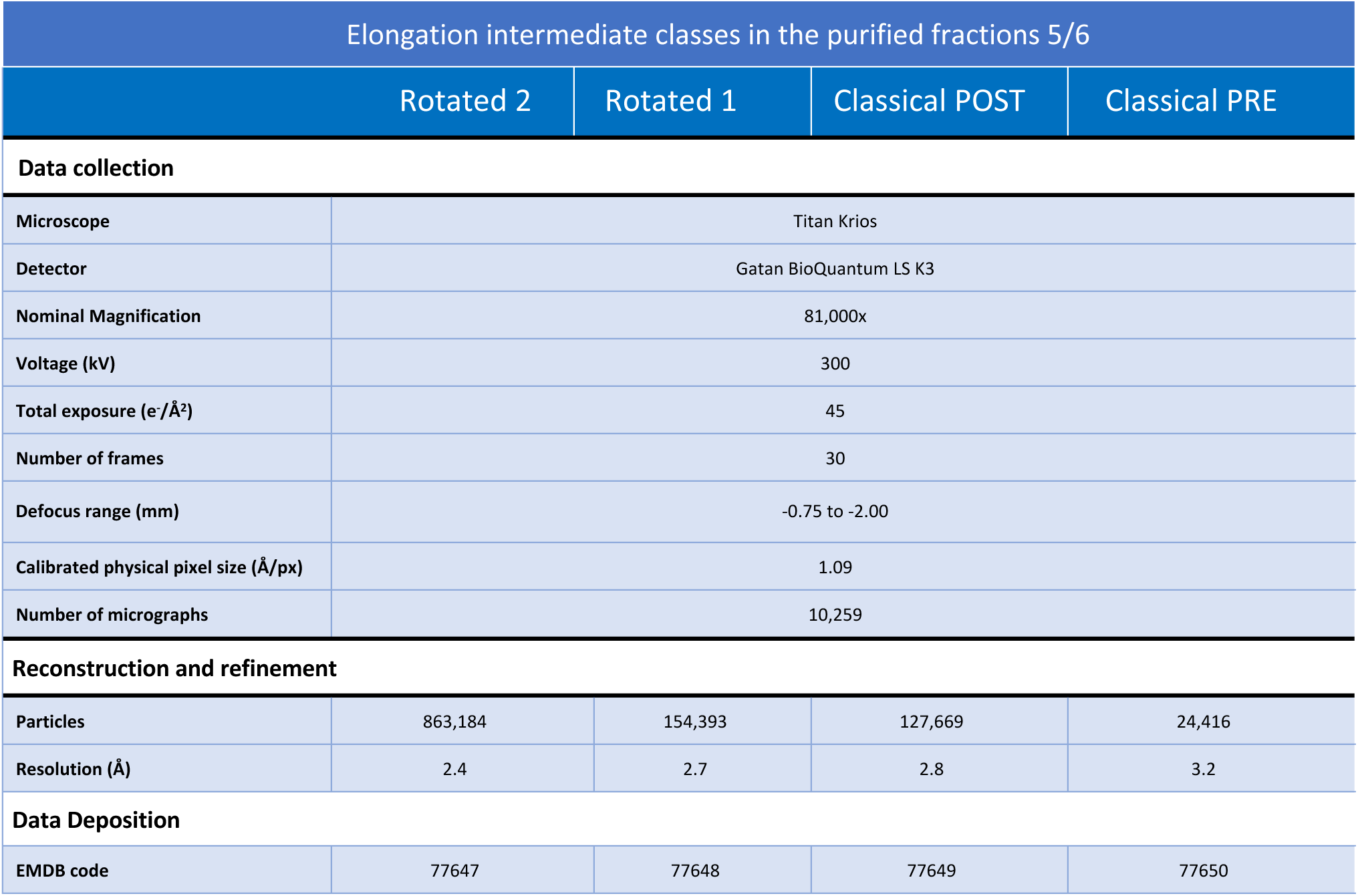
Cryo-EM analysis of the elongation intermediates in the purified fractions 5/6: data acquisition parameters and map statistics.

**Supplementary Table S4.**
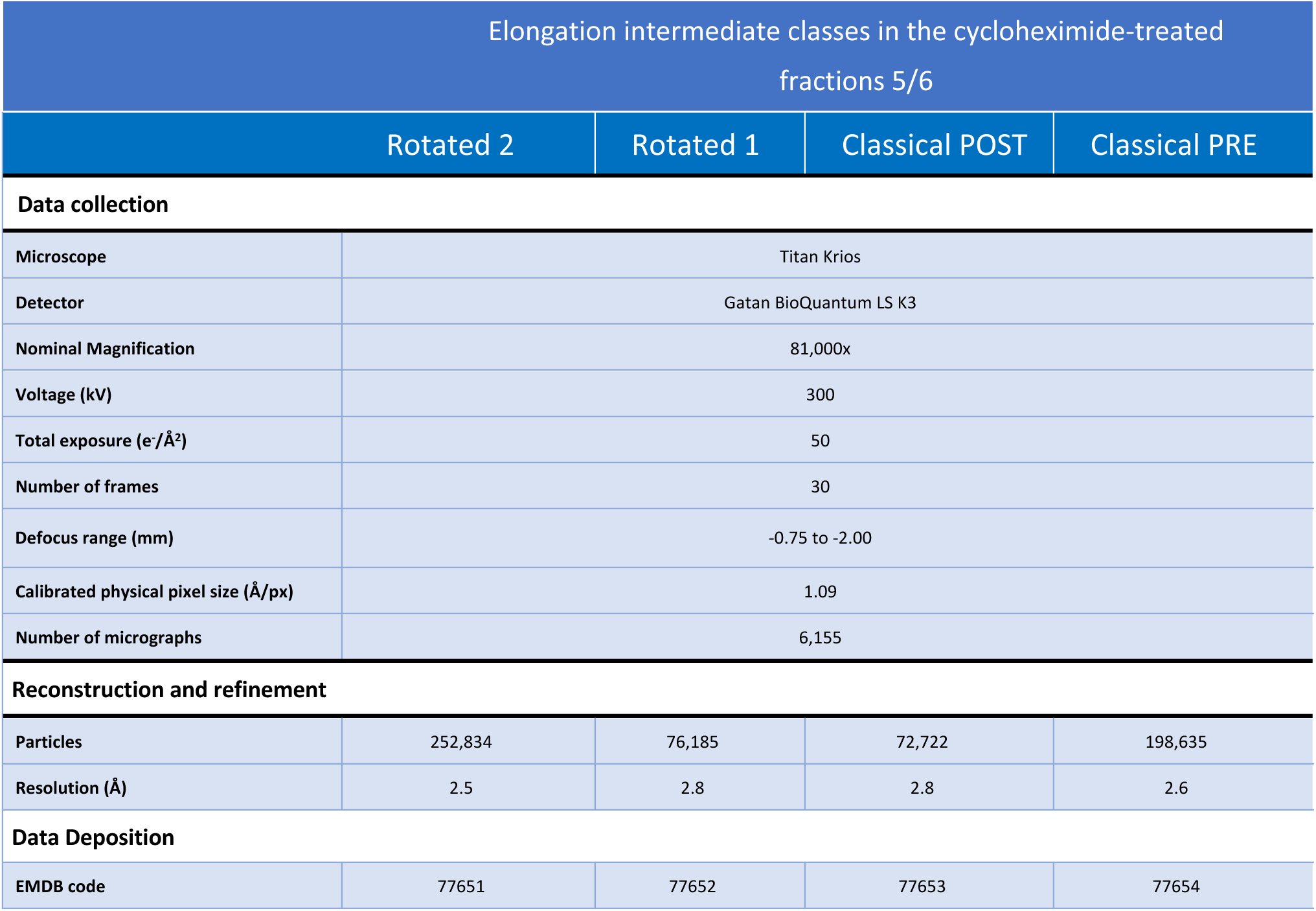
Cryo-EM analysis of the elongation intermediates in the purified cycloheximide-treated fractions 5/6: data acquisition parameters and map statistics.

**Supplementary Table S5.**
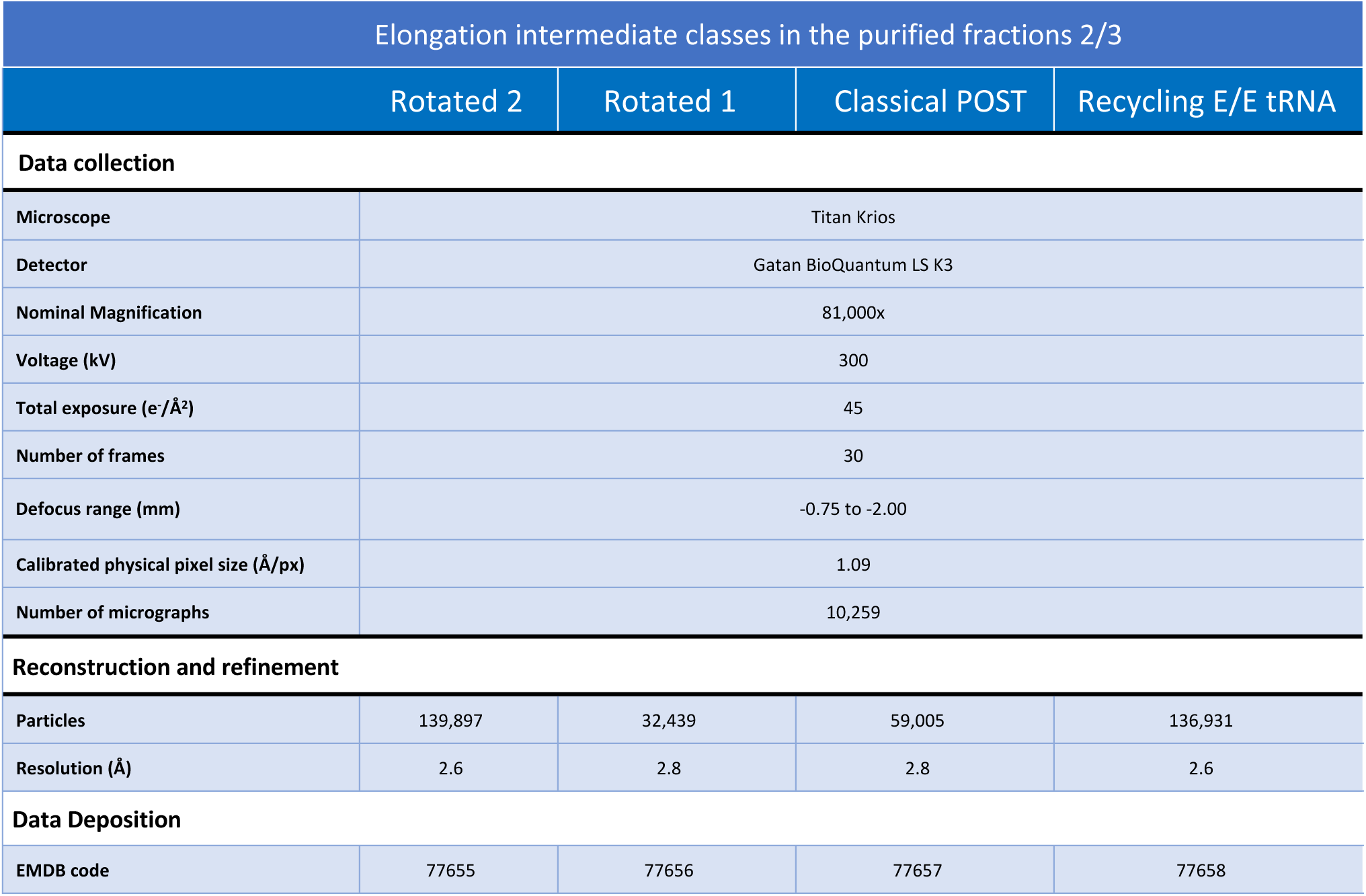
Cryo-EM analysis of the elongation intermediates in the purified fractions 2/3: data acquisition parameters and map statistics.

**Supplementary Table S6.**
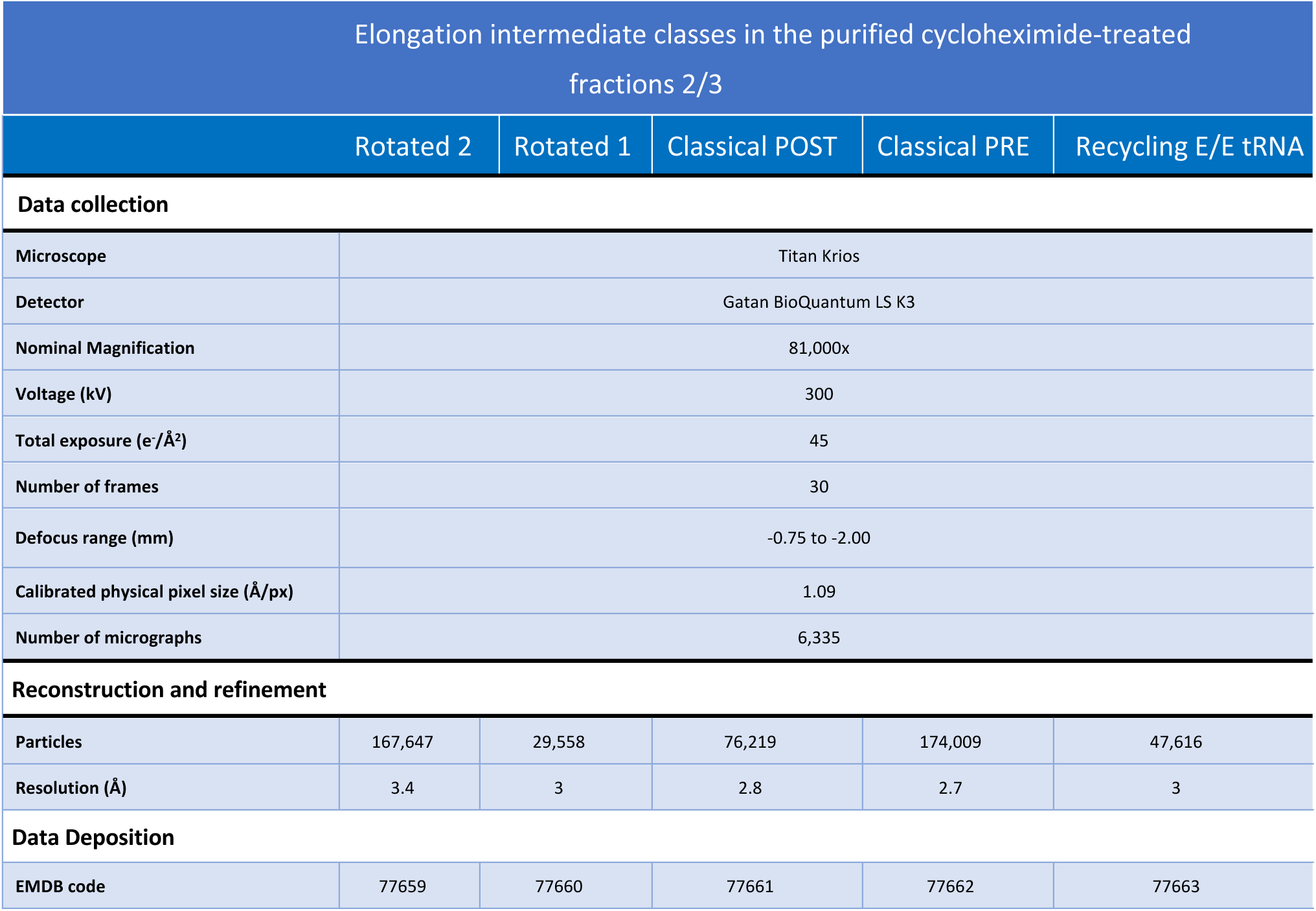
Cryo-EM analysis of the elongation intermediates in the purified cycloheximide-treated fractions 2/3: data acquisition parameters and map statistics.

| Elongation intermediate classes in the purified cycloheximide-treated fractions 2/3 |  |  |  |  |  |
| --- | --- | --- | --- | --- | --- |
|  | Rotated 2 | Rotated 1 | Classical POST | Classical PRE | Recycling E/E tRNA |
| Data collection |  |  |  |  |  |
| Microscope | Titan Krios |  |  |  |  |
| Detector | Gatan BioQuantum LS K3 |  |  |  |  |
| Nominal Magnification | 81,000x |  |  |  |  |
| Voltage (kV) | 300 |  |  |  |  |
| Total exposure (e-/Å²) | 45 |  |  |  |  |
| Number of frames | 30 |  |  |  |  |
| Defocus range (mm) | -0.75 to -2.00 |  |  |  |  |
| Calibrated physical pixel size (Å/px) | 1.09 |  |  |  |  |
| Number of micrographs | 6,335 |  |  |  |  |
| Reconstruction and refinement |  |  |  |  |  |
| Particles | 167,647 | 29,558 | 76,219 | 174,009 | 47,616 |
| Resolution (Å) | 3.4 | 3 | 2.8 | 2.7 | 3 |
| Data Deposition |  |  |  |  |  |
| EMDB code | 77659 | 77660 | 77661 | 77662 | 77663 |

**Supplementary Table S7.**
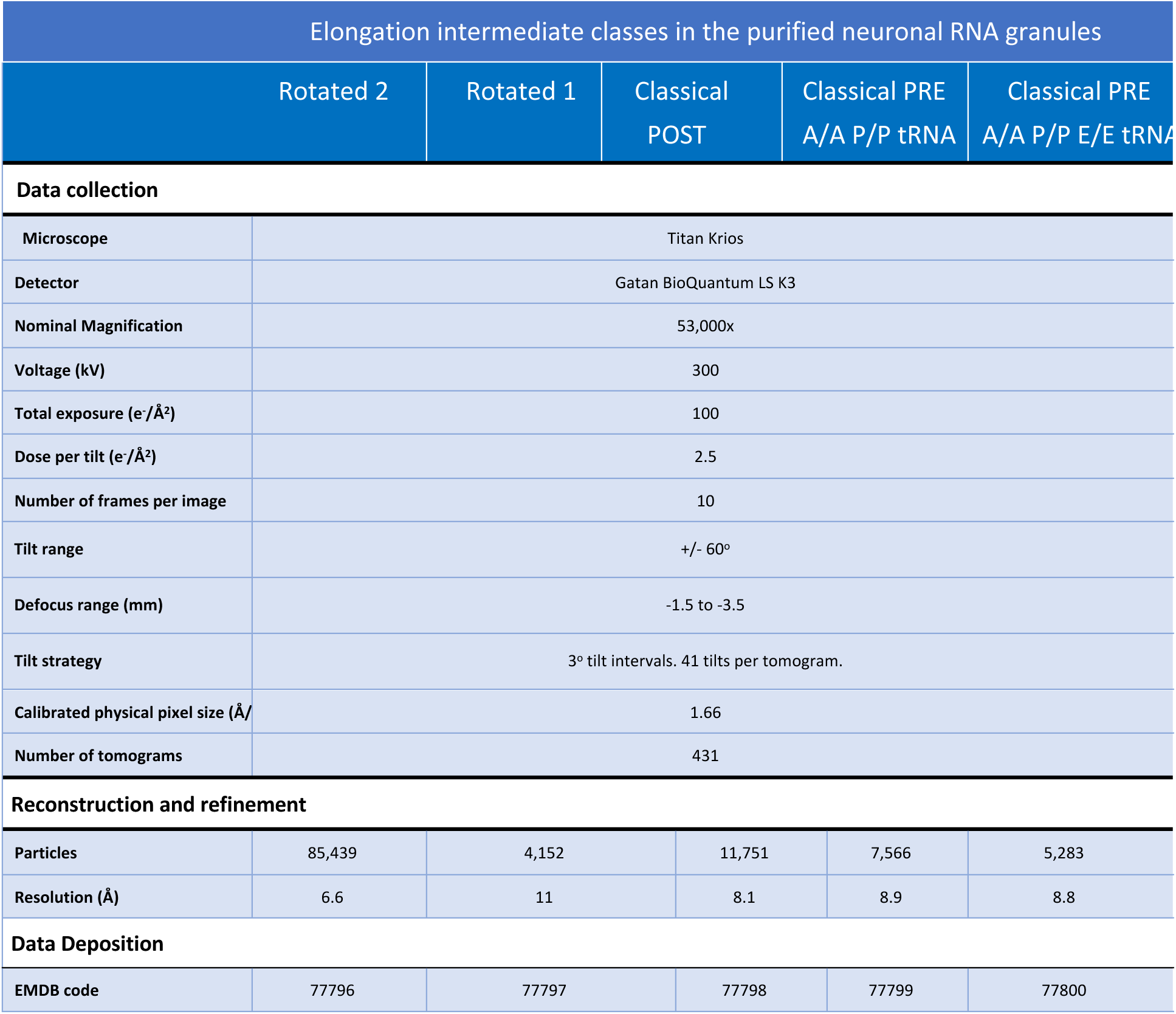
Cryo-ET analysis of the elongation intermediates in the purified dense ribosome clusters from the pellet fraction: data acquisition parameters and map statistics.

